# Oligomerization of PRMT1 Regulated RNA-Binding Protein Cascade Promotes Pancreatic Ductal Adenocarcinoma

**DOI:** 10.1101/2024.12.10.627716

**Authors:** Yanxia Ru, Xinyi Zhou, Xijiao Wang, Wenxin Sun, Yaohui He, Guosheng Hu, Wenjuan Li, Die Hu, Meizhi Jiang, Zhiming Su, Fengfeng Niu, Gang Chen, Jinzhang Zeng, Sen-Fang Sui, Wen Liu, Yaowang Li, Siming Chen

## Abstract

PRMT1 is the predominant type I protein arginine methyltransferase responsible for generating monomethylarginine and asymmetric dimethylarginine (MMA and ADMA) in various protein substrates. It regulates numerous biological processes, including RNA metabolism, mRNA splicing, DNA damage repair, and chromatin dynamics. The crystal structure of rat PRMT1 has been previously reported as a homodimer; however, higher-order oligomeric species of human PRMT1 have been observed in vivo, and their structural basis remains elusive. In this study, we present the cryo-EM structure of human PRMT1 in its oligomeric form, revealing novel interfaces crucial for PRMT1 self-assembly and normal function. PRMT1 shows a strong preference for intrinsically disordered RGG/RG motifs, which are commonly found in RNA-binding proteins (RBPs), highlighting its critical role in regulating RNA metabolism. In vitro studies indicate that disrupting PRMT1 oligomerization impairs its binding affinity for RGG motif substrates, thereby reducing arginine methylation levels on these substrates. This finding emphasizes the essential role of the oligomeric state of PRMT1 in its function with RGG motif-containing RBPs. In vivo, the loss of PRMT1 oligomerization leads to decreased global ADMA levels in pancreatic ductal adenocarcinoma (PDAC) cells and inhibits PDAC tumor growth. Collectively, we propose that PRMT1 oligomerization, rather than mere dimerization, is sufficient for PRMT1-driven PDAC tumor growth, offering novel insights into the potential therapeutic targeting of PRMT1 oligomeric forms in PDAC.

**Graphical abstract:** 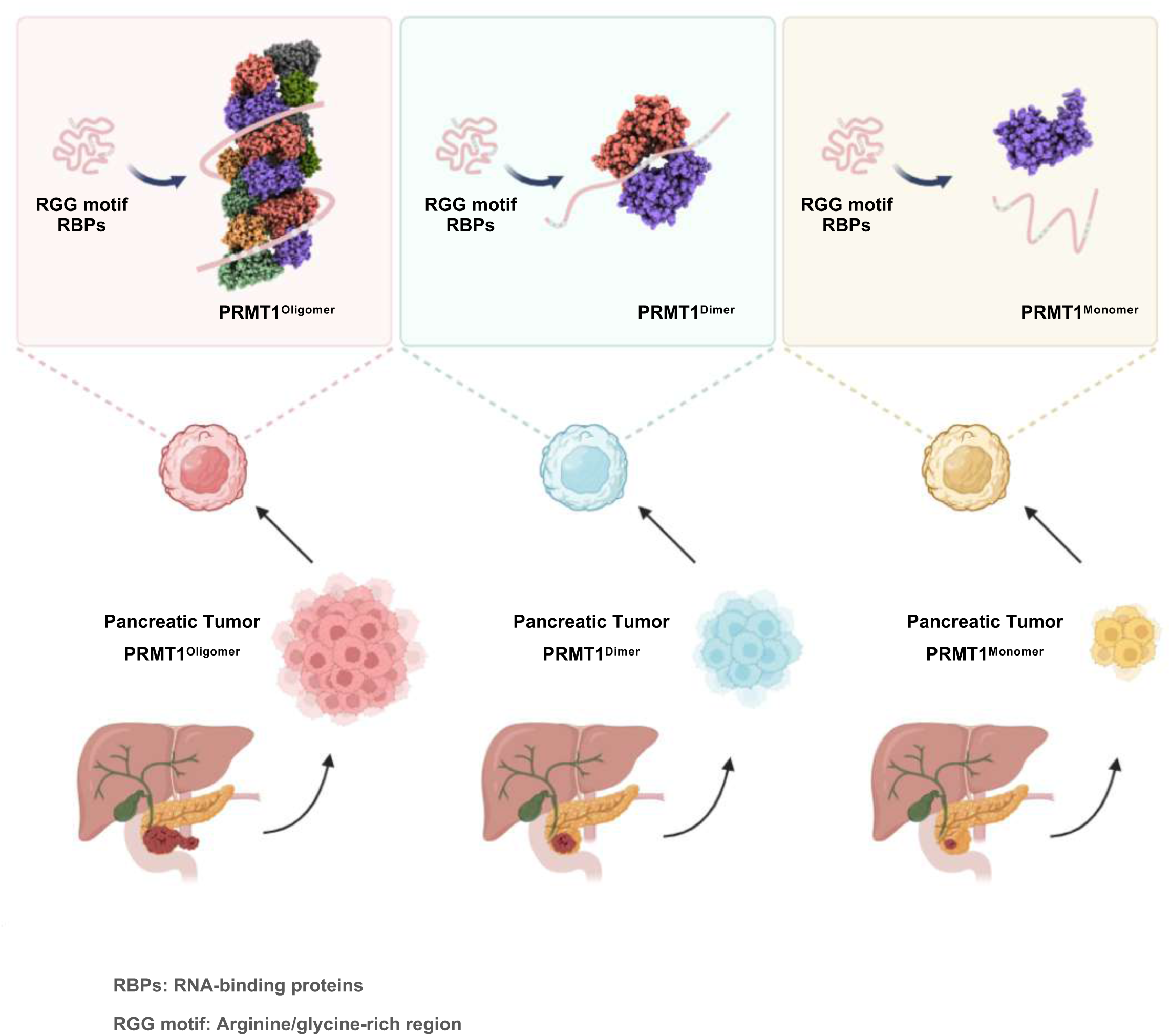

## Introduction

Protein arginine methylation is a ubiquitous post-translational modification in eukaryotic cells, catalyzed by protein arginine methyltransferases (PRMTs)^1^. Currently, nine PRMT members have been identified in humans, which are classified into three distinct types based on their final methyl-arginine products. Type I PRMTs, including PRMT1, PRMT3, PRMT4, PRMT6, and PRMT8, facilitate the formation of asymmetric dimethylarginine (ADMA). In contrast, type II PRMTs (PRMT5 and PRMT9) catalyze the production of symmetric dimethylarginine (SDMA), while type III PRMT (PRMT7) is exclusively responsible for the generation of monomethylarginine (MMA), which serves as an intermediate in both types I and II PRMT reactions^1,2^. PRMTs transfer a methyl group from S-adenosylmethionine (SAM) to their histone or non-histone substrates, thereby modifying their stability, localization, or activity^3^. These modifications significantly influence various cellular processes, including RNA metabolism, DNA repair, chromatin dynamics, and liquid-liquid phase separation (LLPS)^1,3–7^. Therefore, it is not surprising that the expression and activity of PRMTs are tightly regulated, and aberrant arginine methylation is frequently linked to various health disorders, including cardiovascular, neurodegenerative, and cancer-related diseases^6,8^.

PRMT1 is the major type I protein arginine methyltransferase in mammalian cells, accounting for 85% of cellular PRMT activity^9^. Knockout of PRMT1 in mice results in embryonic lethality, underscoring its critical role in embryogenesis and development^10^. Furthermore, aberrant expression of PRMT1 has been documented in several cancer types, including breast, colorectal, and pancreatic cancers^11–16^. Inhibiting PRMT1 activity effectively restricts tumor cell growth, positioning it as an attractive target for therapeutic intervention^6,8^. The crystal structure of rat PRMT1 reveals that it adopts a head-to-tail homodimeric architecture, which is facilitated through its dimerization arm^17^. Mutations or removal of the dimer arm disrupt PRMT1 dimerization, leading to a loss of its activity^17^. These findings underscore the critical importance of the dimeric state for PRMT1’s proper biological functions. Actually, dimerization is a feature conserved across all type I PRMTs and is essential for their normal catalytic activity^18^.

Although it is well recognized that dimerization is required for PRMT1 to perform its biological functions, a series of studies indicate that PRMT1 forms large oligomeric species in vivo^17,19–25^. For instance, either in undifferentiated human primary keratinocytes or murine erythroleukemia (MEL) cells, both endogenous and overexpressed PRMT1 exhibit similar elution behavior to elute in fractions with a molecular mass ranging from 250 to 600 kDa^19,20^. In addition, although the crystal structure of rat PRMT1 has previously been reported as a homodimer, higher-order oligomeric forms of rat PRMT1 are also observed in solution^17^. Collectively, these findings strongly imply that PRMT1 may function in a higher-order oligomeric state in vivo. However, whether these higher oligomeric forms confer distinct biological effects compared to the dimeric form remains unclear.

Motifs rich in arginines and glycines were discovered decades ago and were termed the RGG/RG motifs, which are commonly found in RNA-binding proteins (RBPs)^26^. More than 1,000 human proteins contain the RGG/RG motifs, and these proteins regulate numerous biological processes such as RNA metabolism, mRNA splicing, DNA damage repair, mRNA translation, signal transduction, and LLPS^26–29^. It is now well accepted that arginines within RGG/RG motifs are preferred sites for methylation by PRMTs, in which arginine methylation affects direct interactions of RGG motif proteins with RNAs or proteins, their participation in LLPS, and protein localization^7,26,30,31^. Nonetheless, How PRMT1 recognizes its specific substrates, and to what degree this recognition is regulated by its oligomeric state, remain elusive. Given that multiple RGG/RG repeats are frequently observed in RNA-binding proteins (RBPs), such as the C-terminus of nucleolin (<u>RGG</u>GFGG<u>RGG</u>FGD<u>RGGRGG</u>G<u>RGG</u>) harbors five RGG/RG repeats (underlined), and considering that PRMT1 may function in larger oligomers than dimers in vivo, we hypothesize that PRMT1 utilizes its oligomeric state to recognize RGG motif substrates, particularly those with multiple RGG/RG repeats, through multivalent interactions, thereby enhancing binding affinity and selectivity.

Here, we present the cryo-electron microscopy (cryo-EM) structure of human PRMT1, which adopts a homo-oligomeric architecture featuring a novel helical assembly. Biochemical assays and mutagenesis analyses, combined with size exclusion chromatography (SEC), reveal new interfaces that are critical for PRMT1 self-assembly and normal function. Disrupting PRMT1 oligomerization significantly weakens its binding affinity for substrates containing RGG motifs and reduces the levels of ADMA modifications on these substrates. Furthermore, we demonstrate that the oligomeric state is essential for PRMT1’s role in facilitating the proliferation of pancreatic ductal adenocarcinoma (PDAC) cells. The loss of PRMT1 oligomerization results in decreased global ADMA levels and suppresses PDAC tumor growth. Mechanistically, our transcriptomic and proteomic data indicate that disrupting PRMT1 oligomerization impairs RNA metabolism by reducing its binding affinity for several RNA-binding proteins (RBPs) containing RGG motifs. This finding is further supported previous studies that highlighted the PRMT1-dependent regulation of RNA metabolism in sustaining pancreatic ductal adenocarcinoma^14,32^. Overall, our findings provide structural insights into the oligomeric scaffold of PRMT1, facilitating its recognition of substrates, particularly those RBPs containing multiple RGG repeats, through multivalent interactions, and offer novel perspectives on the potential therapeutic targeting of PRMT1 oligomeric forms in PDAC.

## Results

### Cryo-EM Structure of the Human PRMT1 Helical Polymer

To determine whether PRMT1 can form higher-order oligomers in solution, we subjected purified recombinant human PRMT1 protein to gel filtration chromatography. The PRMT1 protein eluted as a broad peak with a molecular weight ranging from 200 to 440 kDa on a Superose^TM^ 6 Increase 10/300 GL column, indicating its self-association into oligomeric species under the gel filtration buffer conditions (Supplementary Fig. 1A). This observation is consistent with previous findings regarding human PRMT1^17^. To further elucidate the properties of PRMT1 self-association in vitro, we assessed the concentration-dependent behavior of PRMT1 using negative-stain electron microscopy (Fig. 1A). The negative-stain images revealed that PRMT1 can self-assemble into helical polymers in a concentration-dependent manner. The tendency of PRMT1 to form higher-order oligomers increased when the protein concentration exceeded 0.1 mg/ml, while the formation of helical polymers decreased at lower concentrations, specifically below 0.1 mg/ml. However, short helical polymers emerged at 0.2 mg/ml, with their length increasing at concentrations greater than 0.4 mg/ml. Overall, the negative-stain images indicated that the formation of PRMT1 helical polymers is highly dependent on concentration.

**Figure 1.**
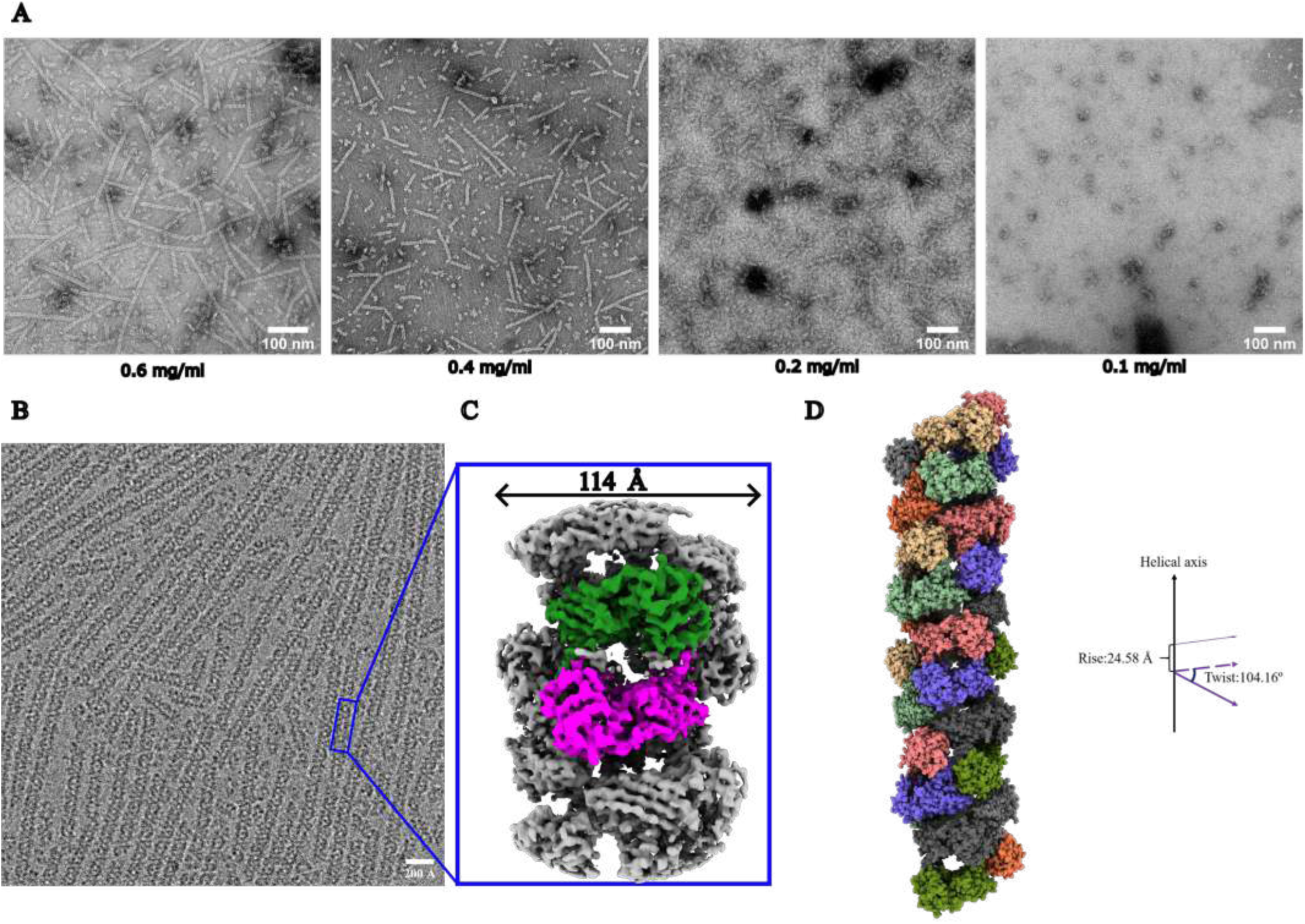
Concentration-dependent PRMT1 helical polymer formation and its helical architecture. (A) The negative stain data of four different concentrations of PRMT1, ranging from 0.6 to 0.1 mg/ml. (B) A cryo-electron microscopy (cryo-EM) micrograph of 0.6 mg/ml PRMT1 was shown, in which all PRMT1 monomers assembled into helical polymers. (C) We solved the helical polymer structure at 3.68 Å, and the PRMT1 colored green and magenta are monomers; the helical polymer diameter is about 114 Å, and its basic unit is a PRMT1 dimer. (D) PRMT1 dimers are assembled into helical polymer one by one at the position after rotating around the helical axis by 104.16° and moving along the direction of the helical axis 24.58 Å distance simultaneously.

Next, we aimed to determine the experimental structure of PRMT1 oligomers using cryo-electron microscopy (cryo-EM) to address existing knowledge gaps between the structural and functional aspects of human PRMT1. Cryo-electron microscopy (cryo-EM) images collected using a Titan Krios microscope (see Table 1 for collection parameters) revealed linear oligomers of varying lengths (Fig. 1B). One representative micrograph indicates that, although some polymers lacked orderly assembly and exhibited heterogeneity, the PRMT1 helical polymers were generally well-distributed (Fig. 1B). Therefore, homogeneous and highly ordered helical polymers were further isolated through classification. Additionally, helical symmetry, a key parameter in helical reconstruction, was estimated using a program developed by the author, which is available for download from GitHub (GitHub link). The cryo-EM map of PRMT1 was reconstructed to a resolution of 3.68 Å (Supplementary Fig. 2). The local resolution is sufficient to determine that the repeating unit of the PRMT1 oligomer is a dimer of PRMT1 (colored in green and magenta), with an approximate diameter of 114 Å (Fig. 1C). A significant number of PRMT1 proteins formed dimers, which subsequently assembled into helical polymer structures following helical symmetry (Fig. 1D). The symmetry positions of the dimers were calculated.

### Multiple Types of Contacts Stabilize the PRMT1 Dimer

As illustrated in Fig. 2A, the identified PRMT1 dimer model serves as a repeating unit within the PRMT1 oligomer. The interface between the two monomers can be directly visualized from the interior perspective of the helical polymer. These interfaces are consistent for each pair of PRMT1 monomers (blue to magenta and magenta to blue in Fig. 2A). PRMT1 adopts a head-to-tail homodimeric orientation, where the head of one monomer contacts the tail of another. Specifically, a loop from one monomer is inserted into the hydrophobic pocket of the other, with amino acid residues W215, W216, V219, Y220, F222, D223, and M224 from the loop interacting with the hydrophobic pocket of the adjacent monomer.

**Figure 2.**
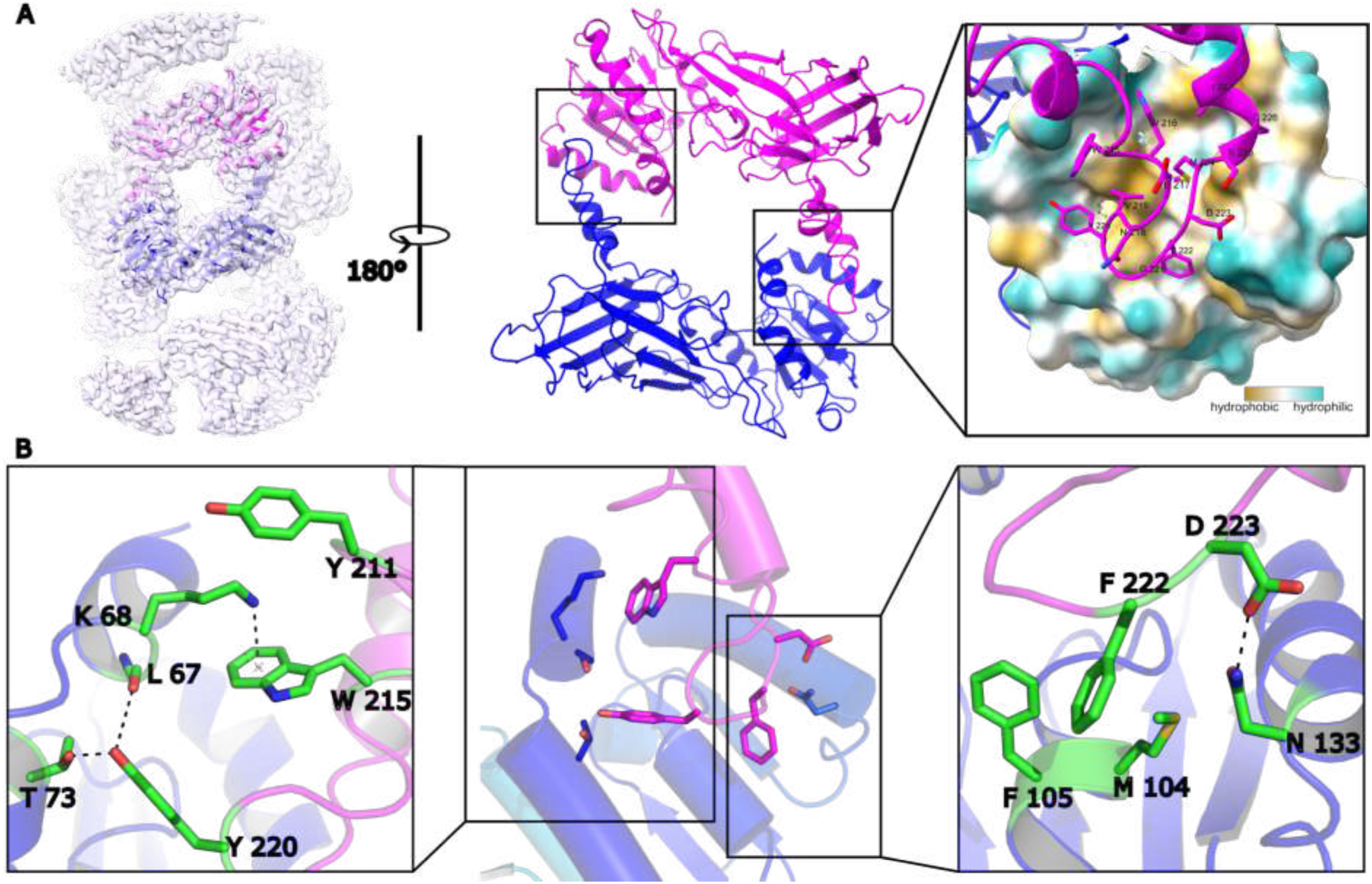
The formation of PRMT1 dimer. (A) The head-tail connection formed a dimer, making it easy to find that a loop in the tail region of one monomer was inserted into the head region of another monomer from the view of the helical polymer interior. The head region of PRMT1 in the interface is a hydrophobic pocket to hold the tail region of another PRMT1. (B) the contacts in the interface were shown as two parts: a cation-pi contact between W 215 and K 68 was found, and a hydrogen bond can be formed between Y220 and L67 (or T73) in the left part; the hydrophobic contact was formed that F222 inserts into the center of M104 and F105 and a hydrogen bond was formed between D223 and N133 in the correct part.

Although the interface between the two monomers is primarily composed of several hydrophobic contacts, electrostatic interactions also contribute to the stabilization of the PRMT1 dimer. Fig. 2B illustrates various types of interactions between amino acid residues at the interface of the two monomers. For example, a cation-π interaction between K68 and W215, combined with a hydrogen bond between L67 (or T73) and Y220, exerts a strong force on the left edge of the hydrophobic pocket. On the right side, a typical hydrophobic contact is observed, with F222 positioned between M104 and F105. Additionally, another hydrogen bond forms between N133 and D223 at the right edge of the hydrophobic pocket. In summary, the cation-π and hydrogen bond interactions on the left, the hydrophobic contacts in the center, and the additional hydrogen bond on the right collectively stabilize the PRMT1 dimer interface, with W215, Y220, and F222 serving as key residues in this stabilization.

### A Unique Pattern of Contacts Is Critical to Assemble PRMT1 Helical Polymer

The formation of helical polymers is dependent on the connections between dimers, which serve as the repeating units of the PRMT1 oligomer. The positions of the three dimers (A, B, and C) are illustrated in Fig. 3A. Two identical interfaces (interfaces 1 and 2) exist between dimers A and B, alongside a weaker contact between dimers A and C (interface 3). In interface 3, T327 interacts with T327, while Y280 from dimer A (shown in green) is inserted into the electrostatic pocket of dimer B (shown in blue), and H296 from dimer B is inserted into the electrostatic pocket of dimer A (Fig. 3A). These residues are depicted in greater detail in Fig. 3B, where Y322 forms hydrogen bonds with F81, Y280 forms bonds with H82 (or N83), and N278 interacts with H296. Contacts at these interfaces strongly couple the dimers, promoting their assembly into helical polymers. Notably, the hydrogen bond pairs, N278 with H296 and Y280 with H82 (or N83), along with adjacent residues, create two “U-shaped” pockets that fit into each other in a face-to-face arrangement (Fig. 3C).

**Figure 3.**
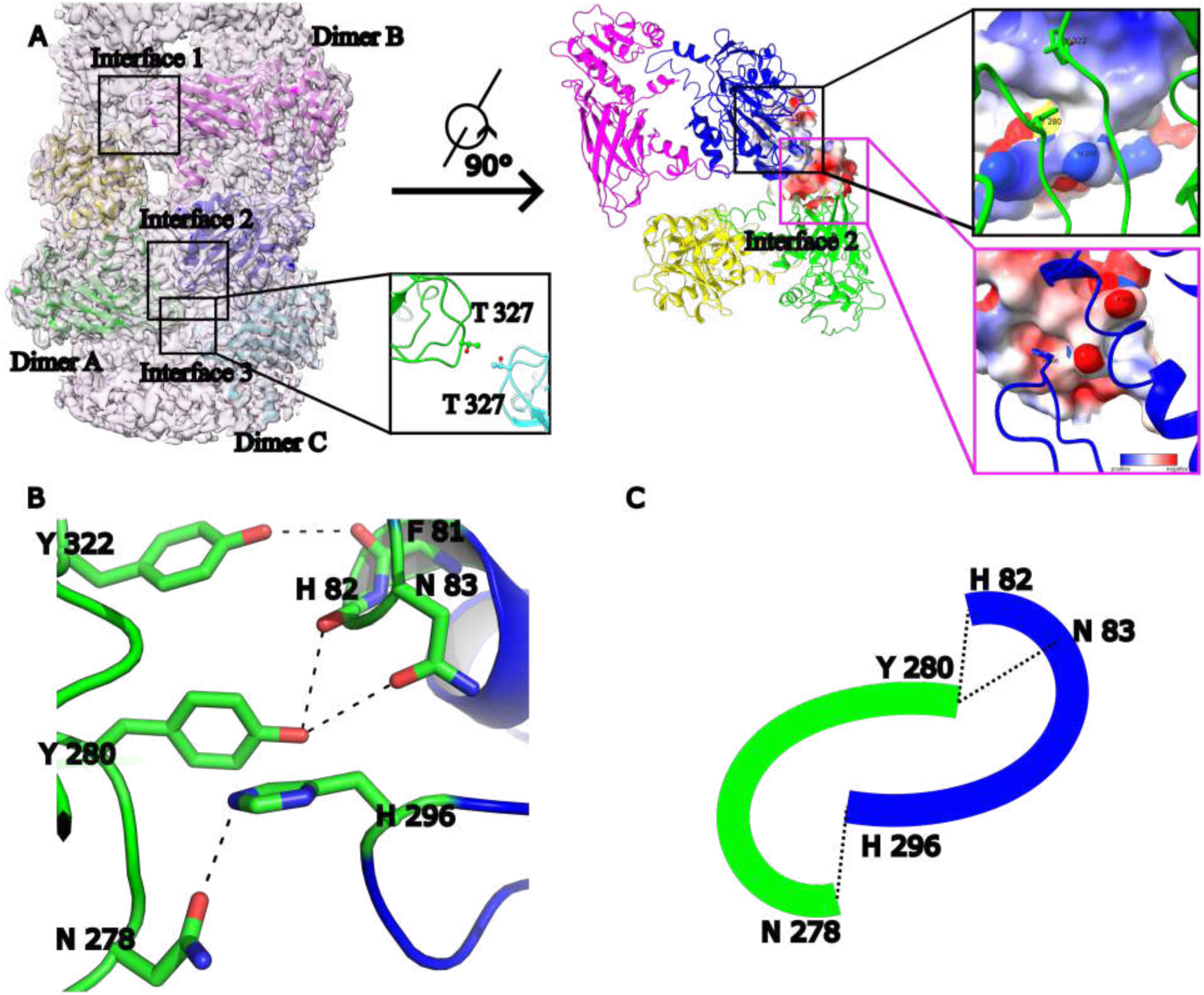
The molecular mechanism of PRMT1 helical polymer. (A) Three dimers connect to form the helical polymer (Dimers A, B, and C). Two identical interfaces (interfaces 1 and 2) between dimers A and B and a weak contact between dimers A and C (interface 3) are shown. T327–T327 on the interface 3. Y280 on the loop from dimer A (green) is inserted into the electrostatic pocket of dimer B (blue). H296 on the loop from dimer B is inserted into the electrostatic pocket of dimer A. (B) The contact in the interface contains three hydrogen bonds, which are N278 and H296, Y280 and H82 (or N 83), Y322 and F81. (C) These contacts form an interesting pattern in two “U-shape” pockets inserted into each other face-to-face.

The current structure allowed us to design various PRMT1 mutations that selectively disrupt PRMT1 oligomers into dimers or monomers in solution. Different types of interfaces are crucial for forming PRMT1 helical polymers. The first interface occurs between PRMT1 monomers, where the residues W215, Y220, and F222 play key roles in stabilizing the PRMT1 dimer (Fig. 2B). The second interface is formed between dimers, in which the residues Y280 and H296, located within the “U-shaped” pockets, are vital for assembling the PRMT1 dimer into helical polymers (Fig. 3D). To confirm the contribution of these amino acid residues to the stabilization of the dimeric or oligomeric state of PRMT1 in solution, we introduced mutations into these residues to generate distinct oligomeric forms of PRMT1. For instance, the mutants PRMT1^W215A/Y220A/F222A^ and PRMT1^Y280A/H296A/T327A^ were generated to produce the monomeric and dimeric states of PRMT1, respectively. To verify the accuracy of the human PRMT1 structure, gel filtration experiments were performed to examine the effects of these amino acid mutations on the oligomerization of the PRMT1. Consistent with our cryo-electron microscopy (cryo-EM) structure of human PRMT1, we observed that purified wild-type PRMT1 eluted as an oligomer on a size exclusion chromatography (SEC) column, as previously reported^17^. Conversely, under the same experimental conditions, PRMT1^W215A/Y220A/F222A^ and PRMT1^Y280A/H296A/T327A^ behaved as a monomer and a dimer, respectively (Supplementary Fig. 3A). Notably, the single mutations Y280A or H296A in PRMT1 sufficiently disrupted the PRMT1 oligomer into a dimer (Supplementary Fig. 3B).

Consistent with the gel filtration results, negative stain images revealed that the PRMT1^W215A/Y220A/F222A^ mutant was unable to assemble into helical polymers and could not form dimers (Supplementary Fig. 3C). Additionally, 2D classification of these negative stain images further demonstrated that the PRMT1^W215A/Y220A/F222A^ mutant exists as a monomer (Supplementary Fig. 3C). However, the negative stain images and additional 2D classification indicated that the PRMT1^Y280A/H296A/T327A^ mutant is incapable of forming helical polymers but retains the dimeric form (Supplementary Fig. 3C).

Finally, a two-tag co-immunoprecipitation analysis was conducted to verify the critical role of the aforementioned amino acid residues in the formation of PRMT1 oligomers (Supplementary Fig. 3D). In this assay, equal amounts of wild-type or mutant FLAG-PRMT1 and MYC-PRMT1 plasmids were co-transfected into HEK293T cells (Supplementary Fig. 3D). Subsequently, nuclear and cytoplasmic extracts were prepared and separately bound to anti-FLAG resin. MYC-PRMT1 binding, via protein oligomerization or dimerization, was assessed using anti-MYC antibodies. Consistently, mutations of W215, Y220, and F222 completely abolished PRMT1 oligomerization and dimerization; however, the Y280, H296, and T327 mutations allowed PRMT1 dimerization to persist while oligomerization was lost (Supplementary Fig. 3E-F). Thus, W215-, Y220-, and F222-mediated dimer stabilization, along with Y280-, H296-, and T327-mediated oligomer stabilization, were validated in the context of full-length PRMT1 protein in HEK293T cells (Supplementary Fig. 3D-F). In subsequent studies, unless otherwise specified, we refer to wild-type PRMT1^WT^ as the oligomeric form of PRMT1 (hereafter referred to as PRMT1^Oligomer^), the PRMT1^Y280A/H296A/T327A^ mutant as the dimeric form of PRMT1 (hereafter referred to as PRMT1^Dimer^), and the PRMT1^W215A/Y220A/F222A^ mutant as the monomeric state of PRMT1 (hereafter referred to as PRMT1^Monomer^) to investigate the effects of different PRMT1 oligomeric states on its biological functions.

### Oligomerization Is Required for Stable Binding of PRMT1 to Substrates Containing RGG Motifs

The arginine- and glycine-rich (RGG/RG) motifs are preferentially methylated by PRMT1, with many previously identified PRMT1 substrates containing multiple tandem RGG motifs^5,6,26,30,31^. Given that oligomers larger than dimers have been observed for PRMT1 in vivo, an efficient mechanism to enhance substrate binding is to increase binding avidity through protein oligomerization^24,33^. In this regard, the binding of PRMT1 to substrates containing multiple tandem RGG motifs may be facilitated by the oligomeric structural platform provided by PRMT1. To validate this hypothesis, we initially examined the impact of distinct oligomeric states on PRMT1 binding to a range of its substrates, including hnRNPA1, hnRNPA2, hnRNPK, METTL14, GAR1, FUS, Nucleolin, and Fibrillarin, all of which possess more than three tandem RGG motifs and are known to be methylated by PRMT1 in multiple cell lines (Fig. 4A)^34–43^. To achieve this objective, we conducted in vitro pull-down assays to compare the binding affinity of PRMT1 for the aforementioned substrates in the context of PRMT1^oligomers^, PRMT1^dimers^, and PRMT1^monomers^. The GST-tagged and FLAG-tagged RGG motif substrates were expressed and purified from *E. coli* for these assays. Coomassie blue staining clearly demonstrated that the oligomeric form of PRMT1 exhibited a significantly higher binding affinity for these RGG substrates compared to its dimeric and monomeric forms (Fig. 4B-H). Domain mapping using a pull-down assay further demonstrated that the C-terminal RGG-rich region of hnRNPA1 is responsible for the stable binding of PRMT1 (Supplementary Fig. 4A-B). These findings collectively support the notion that the avidity effect mediated by oligomers influences PRMT1 binding to its RGG region-containing substrates.

**Figure 4.**
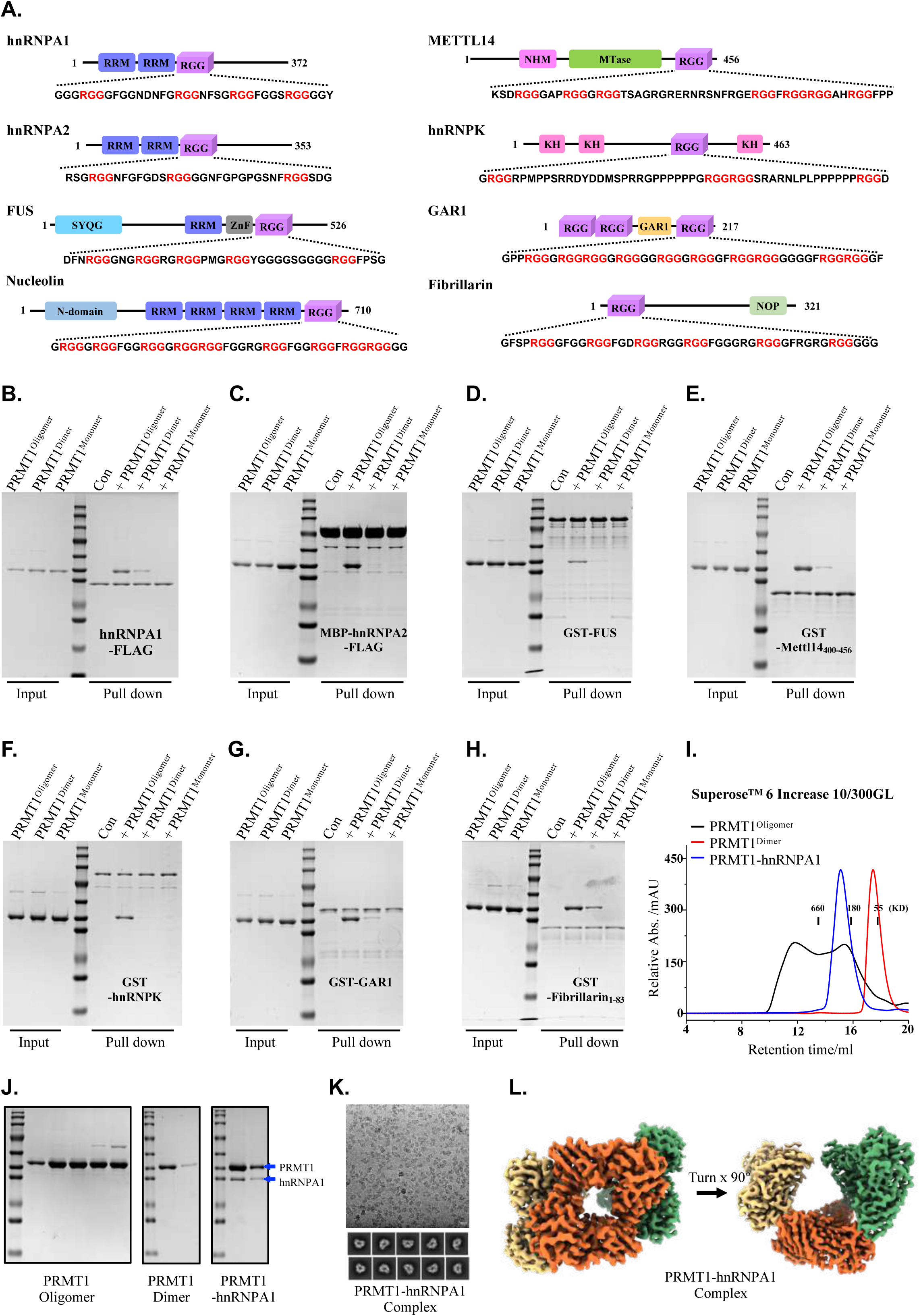
The oligomerization of PRMT1 promotes its binding to substrate proteins enriched with RGG motifs. (A) The domain structure of RGG-rich substrate proteins, including hnRNPA1, hnRNPA2, FUS, Nucleolin, METTL14, hnRNPK, GAR1, and Fibrillarin, as well as the amino acid sequences of their RGG regions, are presented. (B-H) In vitro pull-down experiments were conducted to compare the binding affinities of various proteins, including hnRNPA1-FLAG, MBP-hnRNPA2-FLAG, GST-FUS, GST-METTL14 (400-456), GST-hnRNPK, GST-GAR1, and GST-Fibrillarin (1-83), with different oligomeric states of PRMT1, such as wild-type oligomers, Y280A/H296A/T327A mutant dimers, and W215A/Y220A/F222A mutant monomers. (I-J) The size exclusion chromatography (SEC) elution profile of the PRMT1-hnRNPA1 complex was analyzed. The PRMT1-hnRNPA1 complex, along with wild-type PRMT1 oligomers and Y280A/H296A/T327A mutant PRMT1 dimers, was examined using Superose^TM^ 6 Increase 10/300GL. In this analysis, PRMT1 oligomers and PRMT1 dimers served as controls, and the corresponding gel filtration fraction peaks were collected and analyzed via SDS-PAGE. (K) The cryo-electron microscopy micrograph and the 2D classification results of the PRMT1-hnRNPA1 complex are presented. (L) The 3D reconstruction of the PRMT1-hnRNPA1 complex reveals that it is a hexamer.

We then assessed the extent to which the oligomeric state influences the binding of PRMT1 to substrates containing the RGG region in living cells. FLAG-labeled RGG motif substrates, including hnRNPA1, FUS, hnRNPK, Fibrillarin, GAR1, TAF15, Nucleolin, and RBFOX2, were co-transfected with distinct oligomeric states of MYC-tagged PRMT1, namely MYC-PRMT1^oligomer^, MYC-PRMT1^dimer^, and MYC-PRMT1^monomer^, into HEK293T cells. RGG motif-containing substrates were captured using anti-FLAG resin, and the differential binding of the various oligomeric states of PRMT1 to these substrates was eluted with FLAG peptide and visualized by Western blotting. Consistent with the previously described in vitro pull-down results (Fig. 4B-H), the binding of RGG motif substrates was significantly diminished for PRMT1^monomer^ and PRMT1^dimer^ compared to PRMT1^oligomer^ in HEK293T cells, highlighting emphasizing the crucial role of the avidity effect mediated by PRMT1 oligomerization in RGG motif substrates binding in living cells (Fig. 5A-H).

**Figure 5.**
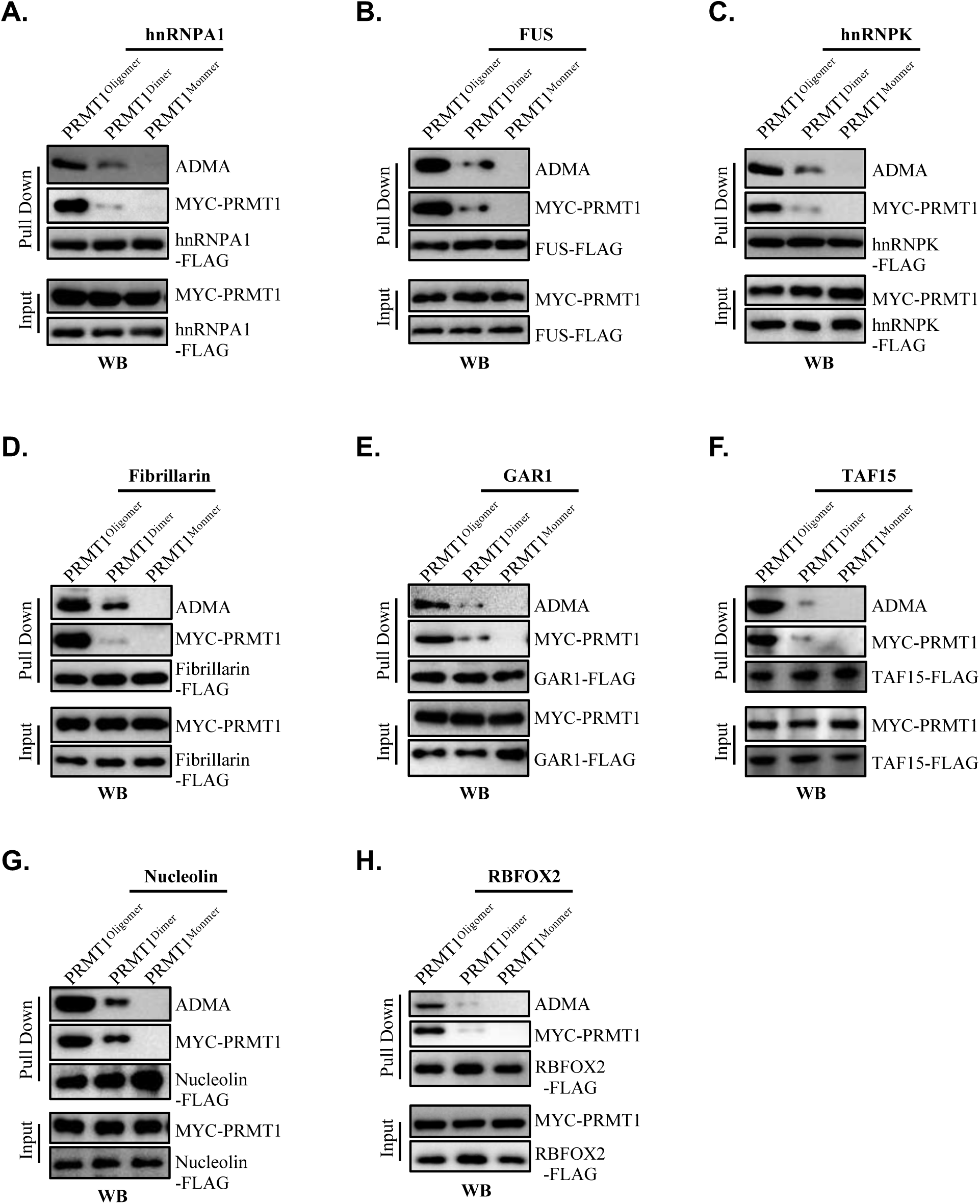
The oligomerization of PRMT1 enhances its binding to and catalysis of substrate proteins enriched with RGG motifs in living cells. (A-H) Co-immunoprecipitation experiments were conducted in HEK 293T cells, where different oligomeric states of MYC-tagged PRMT1—specifically, wild-type PRMT1 oligomers, Y280A/H296A/T327A mutant dimers, and W215A/Y220A/F222A mutant monomers—were co-transfected with FLAG-tagged substrate proteins, including hnRNPA1, FUS, hnRNPK, Fibrillarin, GAR1, TAF15, Nucleolin, and RBFOX2. Cell lysates were prepared and incubated with FLAG affinity resin for binding, followed by elution with FLAG peptide. Anti-MYC signals were utilized to compare the binding affinities between MYC-tagged PRMT1 (oligomers, dimers, or monomers) and the specific substrate proteins. Additionally, the varying anti-ADMA signals in the substrate proteins reflect the different enzymatic activities of the various oligomeric states of MYC-tagged PRMT1.

The mechanism by which PRMT1 specifically methylates certain arginine residues within RGG regions remains unresolved. To address this, we aimed to determine the cryo-electron microscopy (cryo-EM) structure of PRMT1 in complex with RGG-rich substrates to elucidate the structural basis for its recognition of RGG motifs. Fibrillarin and hnRNPA1 were selected as model RGG-rich substrates, and gel filtration experiments were conducted to assess whether PRMT1 could co-elute with either fibrillarin or hnRNPA1 on a gel filtration column. The results indicated that PRMT1 forms stable complexes with both hnRNPA1 and fibrillarin on a size exclusion chromatography (SEC) column (Fig. 4I-J and Supplementary Fig. 4C-D). Finally, single-particle cryo-EM analysis of the PRMT1-hnRNPA1 and PRMT1-fibrillarin complexes confirmed that PRMT1 adopts a hexameric form in both complexes (Fig. 4K-L and Supplementary Fig. 4E-F), further supporting the notion that PRMT1 utilizes its oligomeric form, rather than its dimeric form, to bind RGG-rich substrates. However, we were unable to resolve the structural details of PRMT1 recognition of RGG regions, largely due to the inherent disorder of RGG domains.

To further highlight the role of helical polymers in facilitating PRMT1 binding to RGG-rich substrates, we systematically compared the binding affinities of various PRMT family members for these substrates. PRMT1 and PRMT8 exhibited significantly higher affinities for RGG-rich substrates compared to other PRMT family members, including PRMT2, PRMT3, PRMT5, and PRMT7 (Supplementary Fig. 4G-J). Notably, the previously reported crystal structure of PRMT8 indicates its assembly into helical filaments^44^, which resembles the helical polymer structure of PRMT1 obtained in this study. This finding further emphasizes the importance of the helical polymeric scaffold for PRMTs in their interaction with RGG-rich substrates.

### Oligomerization Significantly Enhances the Enzymatic Activity of PRMT1 toward Substrates Containing RGG Motifs

The binding data presented above raises the question of whether PRMT1 oligomerization enhances its enzymatic activity toward RGG-rich substrates by improving substrate binding. To further explore the relationship between the oligomeric state and the activity of PRMT1, we measured the methyltransferase activity of wild-type PRMT1^oligomer^ and mutant PRMT1^dimer^ against various RGG-rich substrates. Given that dimerization is crucial for AdoMet binding in protein arginine methyltransferases (PRMTs), and that the PRMT1^monomer^ nearly loses its enzymatic activity, we will not further compare the PRMT1^monomer^. The activities of PRMT1^oligomer^ and PRMT1^dimer^ were assessed across a range of concentrations (0.2 µM to 12.8 µM) with saturating concentrations of histone H4, hnRNPA1, and fibrillarin as substrates, respectively, in vitro. Unexpectedly, we found that PRMT1^oligomer^ and PRMT1^dimer^ exhibited nearly identical activity with histone H4, hnRNPA1, and fibrillarin under our experimental conditions (Supplementary Fig. 5A-D). These results clearly demonstrate that both the oligomeric and dimeric forms of PRMT1 are enzymatically active in vitro, supporting the notion that the dimer represents the minimal unit required for PRMT1 methyltransferase activity.

Although there were no significant differences in enzymatic activity between the oligomeric and dimeric forms of PRMT1 toward RGG-rich substrates in vitro, we observed that the assay conditions described above are much simpler than the complex and crowded intracellular environment. Macromolecular crowding in the intracellular milieu is known to influence various protein properties, including protein-protein interactions^45^. We hypothesized that the oligomeric form of PRMT1, due to its higher affinity for RGG-rich substrates, may more effectively overcome binding resistance, thereby promoting the formation of asymmetric dimethylarginine (ADMA) on these substrates in vivo. To test this hypothesis, we compared the different oligomeric states of PRMT1 in catalyzing RGG-rich substrates in HEK293T cells. A series of FLAG-tagged RGG-rich substrates was co-transfected with distinct oligomeric states of MYC-PRMT1 into HEK293T cells. The overexpressed RGG motif substrates, catalyzed by the various oligomeric forms of PRMT1 in living cells, were captured using anti-FLAG resin. Following co-immunoprecipitation, the specific binding of PRMT1 oligomeric states, along with the ADMA-modified RGG substrates, was released by FLAG peptide and visualized by Western blotting. In contrast to the in vitro enzymatic assay results, the oligomeric forms of PRMT1 significantly enhanced enzymatic activity toward RGG-rich substrates. This finding is consistent with the observation that the binding of RGG motif substrates is substantially higher for PRMT1 oligomers compared to PRMT1 monomers and dimers in HEK293T cells, further underscoring the crucial role of PRMT1 oligomerization in binding RGG motif substrates to promote ADMA modification in living cells (Fig. 5A-H).

### Loss of PRMT1 Oligomerization Impairs Global ADMA Levels and Suppresses PDAC Tumor Growth

To investigate how PRMT1 oligomerization affects its biological functions in vivo, it is essential to determine whether higher-order oligomerization occurs under physiological conditions. We selected human pancreatic ductal adenocarcinoma (PDAC) cells based on previous studies that identified PRMT1 as a critical factor for PDAC maintenance^14–16,32^. Additionally, we utilized HeLa cells due to their widespread application in transfection experiments. To exclude the possibility that PRMT1 oligomer formation in living cells results from overexpression, we employed CRISPR-mediated genome editing to knockout endogenous PRMT1 in HeLa cells (Supplementary Fig. 6A-B). Distinct oligomeric states of PRMT1-expressing HeLa cells were generated from the PRMT1 knockout (KO) HeLa cell line. These HeLa cells expressed comparable amounts of PRMT1 oligomers, dimers, and monomers to the endogenous PRMT1 for further investigation (Supplementary Fig. 6C). Similarly, different oligomeric states of PRMT1-expressing PDAC cells were established from PRMT1-depleted human pancreatic cancer cell lines, specifically PANC-1 (Fig. 6A-B).

**Figure 6.**
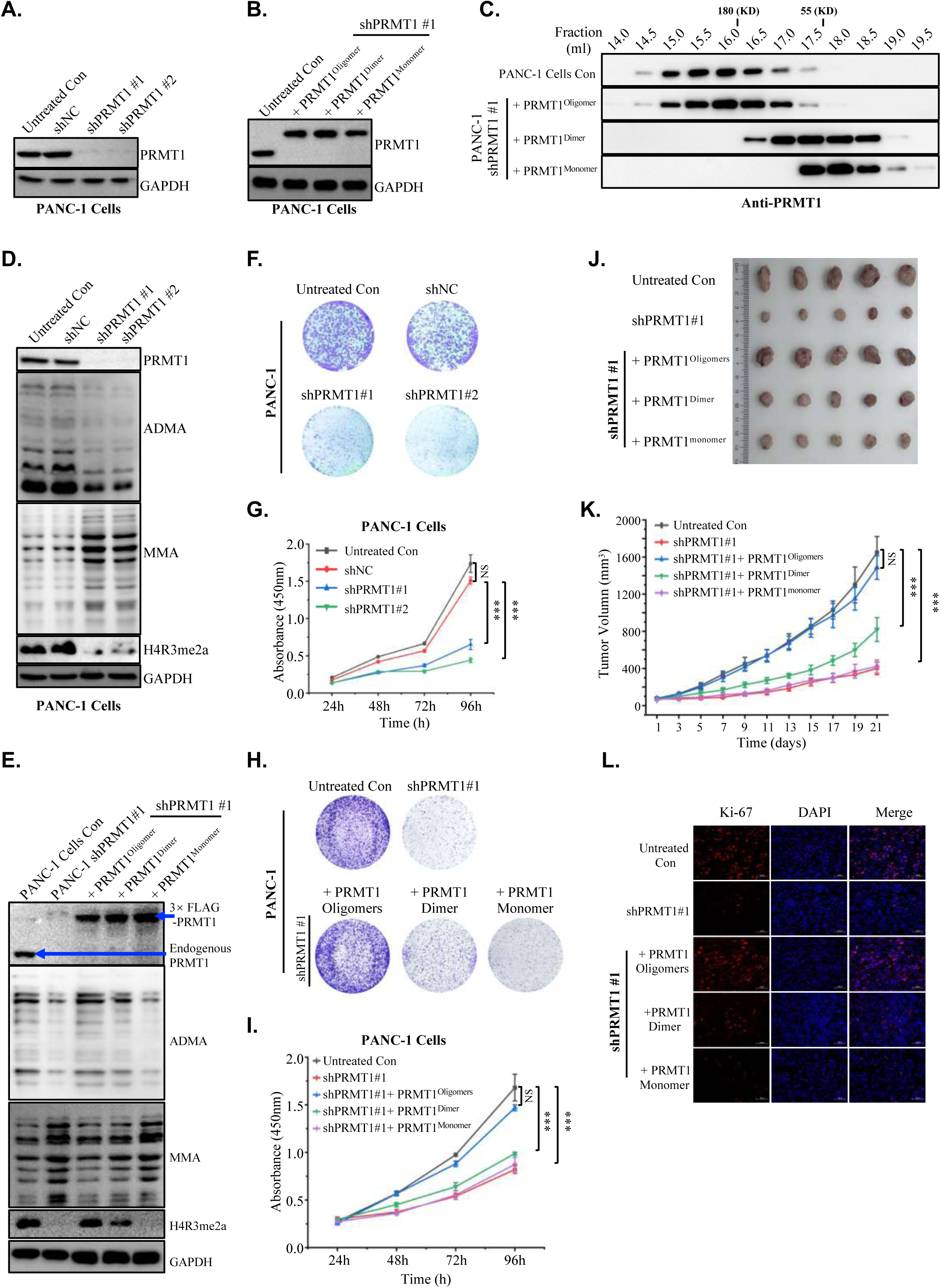
Disruption of PRMT1 oligomerization impairs tumor growth. (A) PANC-1 cells were engineered using two independent PRMT1-targeting shRNAs (shPRMT1#1 and shPRMT1#2) and a non-targeting shRNA (shNC) to generate the corresponding cell lines. Western blot analysis was then performed to assess PRMT1 expression levels, with GAPDH serving as the loading control. (B) PANC-1 shPRMT1#1 cells were transiently transfected with shRNA-resistant PRMT1 oligomers, dimers, and monomers to attain PRMT1 expression levels comparable to those of untreated PANC-1 cells. The results were visualized using Western blot analysis, with GAPDH serving as the loading control. (C) Cell lysates from PANC-1 cells expressing different oligomeric states of PRMT1, as well as untreated PANC-1 cells produced in (B), underwent gel filtration chromatography to verify the endogenous oligomeric state of PRMT1 in PANC-1. Visualization was performed using Western blot analysis. (D) Western blot analysis was conducted to detect the levels of ADMA, MMA, and H4R3me2a in PANC-1 cells following the depletion of PRMT1. GAPDH was included as the loading control. (E) Re-expression of oligomeric PRMT1, dimeric PRMT1, or monomeric PRMT1 in PANC-1 cells following PRMT1 depletion was conducted, and Western blot analysis was performed to measure the levels of ADMA, MMA, and H4R3me2a in the cells, using GAPDH as the loading control. (F-G) Colony formation and proliferation assays were conducted in PANC-1 cells as follows: untreated PANC-1 cells, two independent PRMT1-targeting shRNAs (shPRMT1#1 and shPRMT1#2), and a non-targeting shRNA (shNC), followed by analysis using one-way ANOVA. (H-I) Colony formation and cell proliferation assay results of PANC-1 cells expressing different oligomeric states of PRMT1, along with untreated PANC-1 cells and PRMT1-depleted PANC-1 cells serving as controls, were analyzed using one-way ANOVA. (J) Stable PANC-1 cell lines expressing oligomeric PRMT1, dimeric PRMT1, or monomeric PRMT1 at levels comparable to those of untreated PANC-1 cells were established in PANC-1-shPRMT1#1. These cells were then subcutaneously injected into female BALB/c nude mice for xenograft tumor experiments, and tumors were harvested after 21 days (n=5). (K) Tumor growth curves of the tumors described in (J) were analyzed using two-way ANOVA. (L) Immunofluorescence staining for KI-67 was performed on tumor tissue sections obtained from (J).

Cell lysates from HeLa cells harboring different oligomeric states of PRMT1 were prepared and analyzed using size-exclusion chromatography followed by Western blotting. In untreated HeLa cells, we observed that endogenous human PRMT1 was present at molecular weights exceeding approximately 200 kDa (Supplementary Fig. 6D). A similar elution profile was observed for PRMT1 derived from HeLa cells expressing PRMT1^oligomers^, confirming that the rescue of PRMT1 knockout (KO) HeLa cells with exogenous wild-type PRMT1 preserved an oligomeric state comparable to that of endogenous PRMT1 (Supplementary Fig. 6D). In contrast, PRMT1 from HeLa cells expressing PRMT1 dimers and monomers primarily eluted in a narrow peak corresponding to the dimeric or monomeric state of PRMT1 (Supplementary Fig. 6D). Similar results were also observed in PANC-1 cell lines (Fig. 6C), suggesting that endogenous human PRMT1 exists as a higher-order oligomer in both HeLa cells and the human pancreatic cancer cell line PANC-1.

We subsequently investigated the correlation between the oligomeric states of PRMT1 and its methyltransferase activity in vivo. Recent studies have shown that global inhibition of asymmetric arginine methylation suppresses PDAC tumor growth^14^. Given that oligomerization enhances PRMT1 activity by increasing its binding affinity for RGG-rich substrates, we aimed to determine whether the oligomeric state of PRMT1 is required for the growth of PDAC cells. To validate this hypothesis, we engineered PANC-1 cells using two independent shRNAs targeting PRMT1 or a non-targeting (NT) control shRNA (Fig. 6D). We observed a reduction in ADMA levels and an accumulation of MMA in PRMT1-depleted PANC-1 cells compared to NT shRNA controls (Fig. 6D). Similar results were also observed in MiaPaca-2 (Supplementary Fig. 6E). Ectopic overexpression of an shRNA-resistant PRMT1^oligomer^, but not PRMT1^dimer^ or PRMT1^monomer^ cDNA, restored ADMA and MMA to physiological levels, underscoring the crucial role of the PRMT1 oligomeric state in sustaining its methyltransferase activity in in both PDAC cell lines (Fig. 6E and Supplementary Fig. 7A). Additionally, depletion of PRMT1 significantly inhibited cell proliferation and colony formation in both PDAC cell lines, PANC-1 (Fig. 6F-G) and MiaPaca-2 (Supplementary Fig. 6F-G), indicating a strong dependence of PDAC cells on PRMT1. Moreover, ectopic overexpression of an shRNA-resistant PRMT1^oligomer^, but not PRMT1^dimer^ or PRMT1^monomer^ cDNA, rescued the growth defect in PRMT1-depleted PDAC cells (Fig. 6H-I and Supplementary Fig. 7B-C). Collectively, these data imply that the oligomer, rather than the dimer, is sufficient to promote the growth of PDAC cells such as PANC-1 and MiaPaca-2.

Ultimately, we investigated whether the oligomeric state of PRMT1 is required for maintaining pancreatic ductal adenocarcinoma (PDAC) tumor growth in vivo. PRMT1-depleted PANC-1 cells, rescued with various oligomeric forms, were transplanted into immunocompromised mice, and tumor growth was monitored following tumor establishment. Significant tumor growth inhibition was observed over 21 days in mice with tumors derived from PRMT1-depleted PANC-1 cells compared to control groups (Fig. 6J-K). This growth inhibition could be reversed by reintroducing PRMT1 oligomers, but not PRMT1 dimers or monomers (Fig. 6J-K). These findings suggest that wild-type oligomerization, rather than dimerization, is sufficient to sustain PDAC tumor growth in vivo. Immunofluorescence staining of tumor tissues indicated that the expression levels of the different oligomeric forms of PRMT1 were comparable to endogenous PRMT1 in the control group (Supplementary Fig. 7D). Additionally, immunofluorescence staining for the cell proliferation marker KI-67, along with hematoxylin and eosin (HE) staining, confirmed that the loss of PRMT1 significantly inhibits tumor growth compared to the control group (Fig. 6L and Supplementary Fig. 7E). The reintroduction of PRMT1 oligomers, but not dimers or monomers, reversed tumor growth inhibition (Fig. 6L and Supplementary Fig. 7E). Collectively, these results underscore the critical dependence of established human PDAC xenografts on the oligomeric state of PRMT1 in vivo.

### Disruption of PRMT1 Oligomerization Inhibits PDAC Tumor Growth Through Affecting RNA Metabolism

Among the numerous known RNA-binding proteins (RBPs), those containing RGG motifs rank as the second most prevalent RBPs in the human genome^26,46,47^. RGG motif proteins play critical roles in regulating various facets of RNA processing, including mRNA transport, splicing, and translational regulation^47^. RBPs that possess RGG/RG regions can form high-affinity interactions with RNA^48–50^. Methylation of RGG/RG regions by PRMT1 may influence these interactions, thereby affecting RNA metabolism^51^. PRMT1 has been identified as a critical factor in the maintenance of pancreatic ductal adenocarcinoma (PDAC) due to its fundamental role in RNA metabolism^14,32^. Given that the oligomeric state is essential for PRMT1 to achieve stable binding to RGG-rich substrates and facilitate their methylation in vivo, we investigated whether perturbations in PRMT1 oligomerization, which we have observed to inhibit PDAC tumor growth, are closely linked to alterations in the functions of a subset of RNA-binding proteins containing the RGG motif.

To explore the potential correlation between the disruption of PRMT1 oligomerization and alterations in RNA metabolism, we employed a mass spectrometry-based proteomics approach to identify the differential substrates recognized by various oligomeric states of PRMT1 in PANC-1 cells (Fig. 7A). The distinct oligomeric states of PRMT1 were enriched from PANC-1 cells through immunoaffinity purification using Anti-FLAG resin and subsequently identified via liquid chromatography-tandem mass spectrometry (LC-MS/MS) (Fig. 7A). Analysis of the proteomics results revealed significant changes in peptide abundance among the different oligomeric states of PRMT1, indicating oligomeric state-regulated binding of PRMT1 to substrates (Fig. 7B). To gain further insights into the cellular functions of these substrates, which are differentially recognized by PRMT1 in its various oligomeric states, we performed gene ontology (GO) analysis by mapping the peptides back to their original proteins. We identified 36 proteins out of a total of 619 that were significantly enriched in the oligomeric form of PRMT1 compared to its dimer and monomer forms (Fig. 7B-D). These differentially enriched substrates were primarily associated with nine biological processes in the GO analysis (q < 0.05), including RNA stabilization, RNA splicing, and regulation of the mRNA catabolic process, with RNA stabilization being the most significantly enriched category (q < 0.001; 15 out of 36 RNA stabilization genes enriched) (Fig. 7E). Many differentially bound proteins were RNA-binding proteins (RBPs), including multiple heterogeneous nuclear ribonucleoproteins (hnRNPs) and other RGG motif-containing proteins involved in RNA metabolism (Fig. 7C). This finding is consistent with our in vitro binding results and enhances the confidence in our experimental findings.

**Figure 7.**
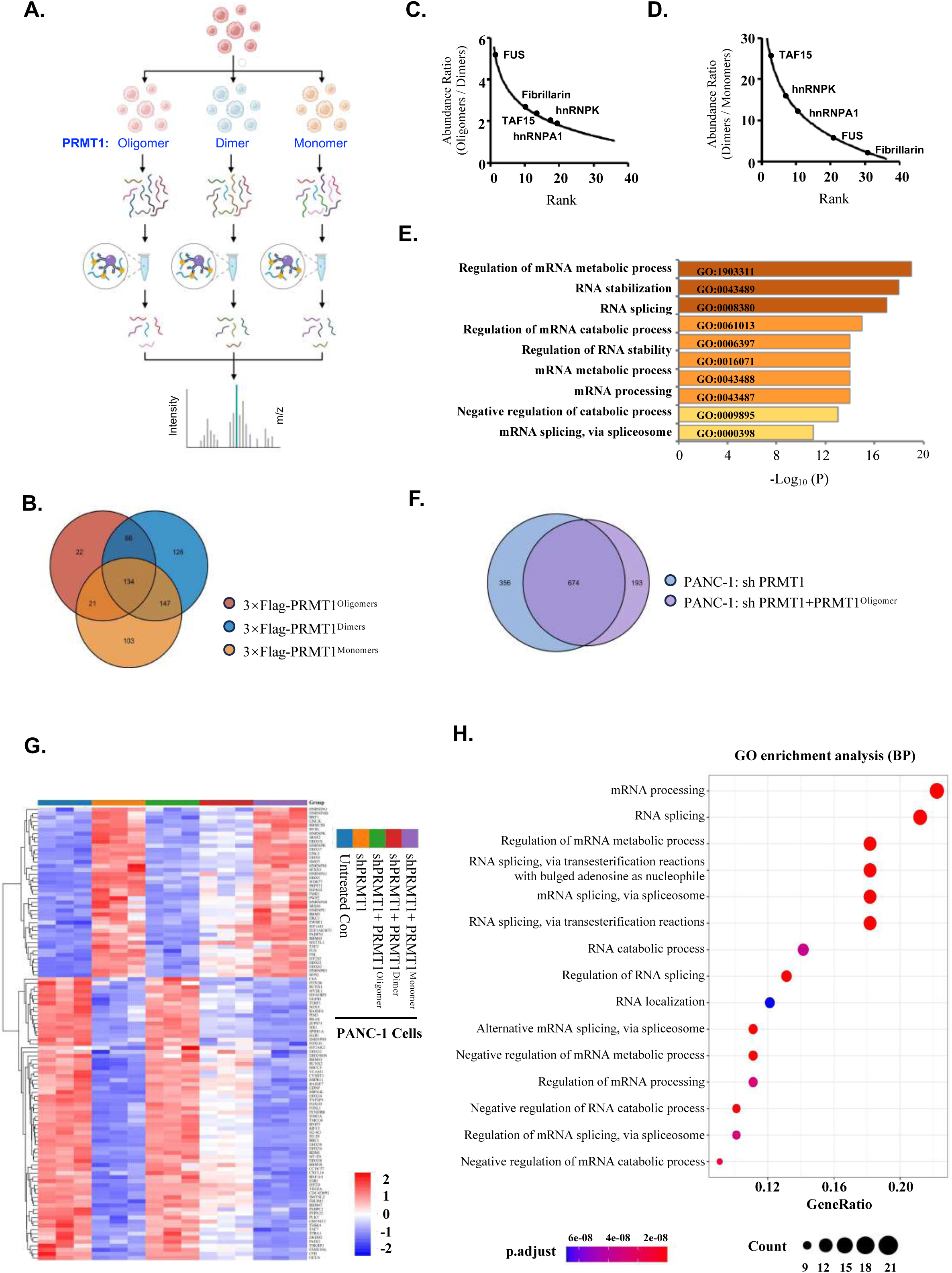
The oligomerization of PRMT1 promotes the growth of pancreatic ductal adenocarcinoma by influencing the cascade of RNA-binding proteins. (A) A flowchart illustrating the experimental procedure for identifying the differential cellular substrates recognized by various oligomeric states of PRMT1 in PANC-1 cells using a mass spectrometry-based proteomics approach is provided. (B) A Venn diagram illustrating the high-affinity substrates associated with PRMT1 oligomers, dimers, and monomers from the experiment described in (A) is presented. (C) Among the 66 substrate proteins identified in the overlap between substrates enriched from PRMT1 oligomers and dimers derived from the Venn diagram in (B), 36 proteins were selected based on an abundance ratio (Oligomers/Dimers) greater than or equal to 2, indicating significant differences in binding between the oligomeric and dimeric states of PRMT1. (D) Abundance ratios of the differential proteins bound by dimers and monomers, as presented in (C). (E) Gene Ontology biological process enrichment analysis of the substrates with varying affinities from (C) and (D) was conducted using Metascape (https://metascape.org), revealing nine enriched gene pathways. (F) A Venn diagram illustrating the overlap between PRMT1 knockdown and the re-expression of PRMT1 oligomers in PRMT1 knockdown cells. (G) A heatmap displaying gene expression levels affected by the different oligomeric states of PRMT1. (H) Gene Ontology biological process enrichment analysis of the relevant genes from (G) was conducted using Hiplot Pro (https://hiplot.com.cn).

Next, we utilized various oligomeric states of PRMT1-expressing PANC-1 cells, alongside untreated PANC-1 cells as a positive control and PRMT1-depleted PANC-1 cells as a negative control, to perform RNA sequencing (RNA-Seq) analysis. To identify the genes regulated by PRMT1, we employed the Venn tool, which identified 674 genes that were significantly altered by PRMT1 depletion and could be rescued by the re-expression of the oligomeric state of PRMT1 in PRMT1-depleted PANC-1 cells (Figure 7F). Subsequently, we compared the transcriptomes of these 674 genes in PANC-1 cells harboring different oligomeric states of PRMT1, including oligomeric, dimeric, and monomeric states, ultimately identifying 112 genes closely correlated with the different oligomeric states of PRMT1, as shown in the heat map (Figure 7G). Gene enrichment analysis of these 112 related genes indicated that they were associated with RNA-related signaling pathways, including “mRNA processing,” “RNA splicing,” and “regulation of mRNA metabolic processes,” which align closely with the gene ontology (GO) analysis derived from the aforementioned LC-MS/MS results (Figure 7E and 7H). Collectively, these results further reinforce the necessity of the oligomeric state for PRMT1’s functionality in pancreatic ductal adenocarcinoma (PDAC) and confirm that the disruption of PRMT1 oligomerization suppresses tumor growth by influencing RNA metabolism.

## Discussion

Several studies have demonstrated that PRMT1 forms oligomers larger than dimers in vitro through the utilization of size-exclusion chromatography (SEC), analytical ultracentrifugation (AUC), and chemical crosslinking^21,24,44,52–55^. However, the biological function of PRMT1 oligomers in vivo has not yet been fully elucidated. Dimerization is a conserved characteristic among all type I PRMTs and is essential for normal catalytic activity^18^. Consequently, dimerization has long been regarded as necessary for the biological functions of PRMT1; yet, it remains uncertain whether intracellular dimerization alone suffices for PRMT1 to fulfill these functions and whether higher-order oligomerization occurs under physiological conditions. Few studies have systematically compared the biological functions of oligomers and dimers in vivo, largely due to a lack of structural data regarding the oligomeric state of human PRMT1. Although the crystal structure of rPRMT1 has been previously reported as a homo-dimer, higher-order oligomers were detected in solution^17^, suggesting that crystallization conditions may have disrupted rPRMT1 oligomerization^17,24^. To date, a homo-oligomeric structure has been identified in Saccharomyces cerevisiae PRMT1 (ScPRMT), which forms a ring-like hexamer^22^; however, ScPRMT1 shares only 50% sequence identity with hPRMT1. Thus, the structural basis for hPRMT1 oligomerization remains unresolved.

In this study, we determined the cryo-electron microscopy (Cryo-EM) structure of human PRMT1 in its oligomeric form. Furthermore, we characterized hPRMT1 using a combination of biochemical assays, mutagenesis experiments, and functional assays. Size-exclusion chromatography revealed that PRMT1 elutes as a broad peak ranging from 200 to 600 kDa, indicating the formation of heterogeneous oligomers. Negative staining further corroborated that PRMT1 assembles into a concentration-dependent helical polymer. We demonstrated that endogenous PRMT1 exists as oligomers larger than dimers under physiological conditions. Analysis of the Cryo-EM structure of hPRMT1 led to the identification of two sets of amino acid residues at different interfaces of the oligomers, which are involved in stabilizing the dimerization and oligomerization of PRMT1. Additionally, we generated monomeric and dimeric PRMT1 mutants for functional studies. Finally, we revealed that the disruption of PRMT1 oligomers into dimers or monomers significantly impairs its binding affinity for a series of RNA-binding proteins (RBPs) containing RGG/RG motifs, thereby reducing arginine methylation levels on these substrates in living cells.

Proteins containing RGG motifs rank as the second most prevalent group of RNA-binding proteins^26,46,47^. These proteins have been shown to undergo liquid-liquid phase separation both in vitro and in vivo^29,56,57^, a process likely regulated by various post-translational modifications, including phosphorylation and methylation^58^. Several studies have demonstrated that PRMT1 methylates RGG substrates to modulate their phase behavior^7,59^. In this study, we found that the oligomeric state of PRMT1 is required to form a stable complex with RGG motif substrates in a homogeneous hexameric form on an SEC column, at least for hnRNPA1 and fibrillarin that we tested. Given that phase separation is highly dependent on the degree of protein oligomerization, our findings provide additional insight into how PRMT1 may regulate the phase separation of RNA-binding proteins containing RGG motifs beyond its previously established enzymatic activity. This adds another layer of regulatory complexity to the process of phase separation of RGG motif-containing RNA-binding proteins regulated by PRMT1. Further investigation is needed to determine the extent to which the phase separation of RGG substrates depends on PRMT1’s enzymatic activity versus their direct binding. In the future, the application of separation-of-function PRMT1 mutations, based on the cryo-EM structure of hPRMT1 reported here, will enable us to gain a more comprehensive understanding of this topic.

In this work, we screened and identified that both PRMT1 and PRMT8 exhibit significantly higher affinity for substrates enriched with RGG motifs compared to other members of the PRMT family. Structural analyses revealed that both the previously reported crystal structure of PRMT8 and the cryo-electron microscopy (cryo-EM) structure of PRMT1, presented in this study, adopt a similar helical polymer conformation. This finding strongly suggests that the helical polymer structural scaffold may represent a predominant conformation for PRMTs when recognizing substrates containing the RGG motif. Our results indicate that the helical polymer may account for PRMT1’s inherent preference for RGG motifs in vivo and in vitro. To gain further insights into their interaction details, we attempted to resolve the cryo-EM structures of the PRMT1-hnRNPA1 and PRMT1-fibrillarin complexes; however, we were unable to elucidate the structural details of hnRNPA1 or fibrillarin within these complexes due to significant disorder present in the RGG regions.

The regulation of RNA metabolism by PRMT1 has been demonstrated as a primary mechanism for sustaining pancreatic ductal adenocarcinoma^14,32^. In this study, we showed that the depletion of PRMT1 significantly inhibits the proliferation of pancreatic cancer cells, including PANC-1 and MiaPaca-2. Notably, the restoration of PRMT1’s oligomeric forms, but not its dimeric or monomeric forms, can reverse this inhibition, indicating that the oligomeric state of PRMT1 is essential for sustaining pancreatic cancer cell proliferation. To elucidate the mechanisms by which PRMT1 supports this proliferation, we established various oligomeric states of PRMT1 expression in PANC-1 cells and employed a mass spectrometry-based proteomics approach to identify the differential substrates recognized by the oligomeric forms of PRMT1. We conducted Gene Ontology (GO) analysis of proteins enriched in the oligomeric form of PRMT1 compared to its dimeric and monomeric forms, revealing that the differentially enriched substrates are RNA-binding proteins rich in RGG motifs, primarily associated with RNA processing. Therefore, the findings of this study suggest that the disruption of PRMT1 oligomerization significantly inhibits the proliferation of pancreatic cancer cells, likely due to disturbance of PRMT1’s binding to RNA-binding proteins enriched with RGG motifs.

The RGG region in RNA-binding proteins serves as a preferential site for arginine methylation by PRMT1, a process that regulates their binding affinity to RNA. Consequently, alterations in the methylation status of RNA-binding proteins catalyzed by PRMT1 can significantly affect aberrant mRNA biogenesis. Our transcriptomic data suggest that the disruption of PRMT1 oligomerization leads to substantial changes in co-transcriptional RNA processing, including splicing alterations. Thus, in combination with the previously described mass spectrometry-based proteomics data, disruption of PRMT1 oligomerization decreases its binding to several RNA-binding proteins, consequently impairing the methylation of arginine residues in the RGG regions of these substrates. The combined transcriptomic and proteomics data support the conclusion that the loss of PRMT1 oligomerization inhibits pancreatic ductal adenocarcinoma growth by affecting the arginine methylation of RNA-binding proteins, thereby disrupting RNA metabolism.

## Author contributions

S.C., W.L., and Y.L. conceptualized the project, while X.W. prepared the samples for cryo-electron microscopy (Cryo-EM) structure analysis. X.Z., X.W., and W.S. designed and conducted the majority of the experiments with assistance from Y.H., G.H., W.L., D.H., and M.J.; Y.R. was responsible for data acquisition; Y.R., Z.S., and F.N. undertook image processing and structure determination. Supervision of the project was provided by S.C., W.L., and Y.L., and all authors contributed to the data analysis, and manuscript preparation.

## Acknowledgements

This work was supported by the National Natural Science Foundation of China (32270638, 32100464), the National Key Research and Development Program of China (2023YFE0118000), the Shenzhen Science and Technology Innovation Commission (JCYJ20230807091204009), the Natural Science Foundation of Fujian (2022J01051), the Fundamental Research Funds for the Central Universities (20720220122), and the Nanqiang Outstanding Young Talents Program from Xiamen University.

## Ethics statement

The animal study was reviewed and approved by Laboratory Animal Welfare and Ethics Committee of Xiamen University.

