## Supplemental data for "Oligomerization of PRMT1 Regulated RNA-Binding Protein Cascade Promotes Pancreatic Ductal Adenocarcinoma"

**This PDF file includes:**

**Materials and Methods**

**Figs. S1 to S7**

**Table S1**

#### STAR ★ METHODS

##### KEY RESOURCES TABLE

| REAGENT or RESOURCE | SOURCE | IDENTIFIER |
|---|---|---|
| **Antibodies** | | |
| Myc-Tag (19C2) mAb | Abmart | Cat No. M20002S |
| Strep-Tag II Antibody | Abmart | Cat No. M40014S |
| Histone H4R3me2a (asymmetric) (pAb) | Active Motif | Cat No. 39006 |
| pan-Mono-Methyl Arginine Motif Rabbit pAb | Abclonal | Cat No. A17984 |
| Histone H4 Antibody | Cell Signaling Technology | Cat No. 2592S |
| Asymmetric Di-Methyl Arginine Motif [adme-R] Rabbit mAb | Cell Signaling Technology | Cat No. 13522S |
| PRMT1 Polyclonal antibody | Proteintech | Cat No. 11279-1-AP |
| Monoclonal ANTI-FLAG antibody | Sigma-Aldrich | Cat No. F1804 |
| GAPDH Monoclonal Antibody | Invitrogen | Cat No. MA1-16757 |
| **Gel filter column** | | |
| Superose6™ Increase 10/300GL | Cytiva | Cat No. 29091596 |
| Superdex™ 200 Increase 10/300GL | Cytiva | Cat No. 28990944 |
| **Recombinant DNA** | | |
| pET-28a-SUMO-StrepI-Tev-PRMT1 (32-371) | This Paper | N/A |
| pET-28a-SUMO-StrepI-Tev-PRMT1 (32-371) Y280A/H296A/T327A | This Paper | N/A |
| pET-28a-SUMO-StrepI-Tev-PRMT1 (32-371) W215A/Y220A/F222A | This Paper | N/A |
| pET-28a-SUMO-StrepI-Tev-PRMT1 (32-371) Y280A | This Paper | N/A |

| | | |
|---|---|---|
| pET-28a-SUMO-StrepI-Tev-PRMT1 (32-371) H296A | This Paper | N/A |
| pET-28a-SUMO-StrepI-Tev-PRMT1(32-371) F222A | This Paper | N/A |
| pET-28a-SUMO-hnRNPA1 (1-320)-Flag | This Paper | N/A |
| pET-28a-SUMO-hnRNPA1 (186-320)-Flag | This Paper | N/A |
| pET-28a-SUMO-hnRNPA1 (1-196)-Flag | This Paper | N/A |
| pGEX-4T1-Tev-Fibrillarin (1-82) | This Paper | N/A |
| pMAL-C2X-hnRNPA2 (1-341)-Flag | This Paper | N/A |
| pGEX-4T1-Tev-FUS (1-526) | This Paper | N/A |
| pGEX-4T1-Tev-GAR (1-271) | This Paper | N/A |
| pGEX-4T1-Tev-hnRNPK (1-463) | This Paper | N/A |
| pGEX-4T1-Tev-Mettl14 (400-456) | This Paper | N/A |
| pMAL-C2X-Fibrillarin (1-321)-Flag | This Paper | N/A |
| pLKO.1-hygro-shPRMT1#1 | This Paper | N/A |
| pLKO.1-hygro-shPRMT1#2 | This Paper | N/A |
| pLKO.1-hygro-shNC | This Paper | N/A |
| pCDH-EF1-3 × Flag-PRMT1 | This Paper | N/A |
| pCDH-EF1-3 × Flag-PRMT1 (Y280A/H296A/T327A) | This Paper | N/A |
| pCDH-EF1-3 × Flag-PRMT (W215A/Y220A/F222A) | This Paper | N/A |
| pCDH-EF1-Myc-PRMT1 | This Paper | N/A |
| pCDH-EF1-Myc-PRMT1 (Y280A/H296A/T327A) | This Paper | N/A |
| pCDH-EF1-Myc-PRMT (W215A/Y220A/F222A) | This Paper | N/A |
| pCDH-EF1-3 × Flag-GAR1 (1-271) | This Paper | N/A |
| pCDH-EF1-3 × Flag-Fibrillarin (1-321) | This Paper | N/A |

| | | |
|---|---|---|
| pCDH-EF1-3 × Flag-RBFOX2 (1-390) | This Paper | N/A |
| pCDH-EF1-3 × Flag-FUS (1-526) | This Paper | N/A |
| pCDH-EF1-3 × Flag-TAF15 (1-592) | This Paper | N/A |
| pCDH-EF1-3 × Flag-Nucleolin (1-710) | This Paper | N/A |
| pCDH-EF1-3 × Flag-hnRNPA1 (1-320) | This Paper | N/A |
| pCDH-EF1-3 × Flag-hnRNNPK (1-463) | This Paper | N/A |
| lentiCRISPR v2-sgPRMT1 | This Paper | N/A |
| lentiCRISPR v2 | Addgene | Cat No. 52961 |
| psPAX2 | Addgene | Cat No. 12260 |
| pCMV-VSV-G | Addgene | Cat No. 8454 |
| **REAGENT or RESOURCE** | | |
| Polyethylenimine Linear (PEI) MW25000 | YEASEN | Cat No. 40815ES03 |
| Thyroglobulin | Bioss | Cat No. bs-0291P |
| Strep-Tactin®XT 4Flow® high capacity resin | IBA Lifesciences | Cat No. 2-5030-010 |
| Pierce™ Glutathione Agarose | Thermo Scientific™ | Cat No. 16101 |
| FLAG peptides | Sigma | Cat No. F3290 |
| AG RNAex Pro Reagent | Accurate Biotechnology | Cat No. AG21101 |
| Anti-DYKDDDDK G1 Affinity Resin | GenScript | Cat No. L00432-5 |
| Penicillin/Streptomycin | Yeasen | Cat No. 60162ES76 |
| Puromycin (Solution 10 mg/mL) | Yeasen | Cat No.60209ES10 |
| Hygromycin B | MCE | Cat No. HY-B0490 |
| Clarity Western ECL Substrate, 500 mL | Bio-Rad | Cat No.1705061 |
| Cell Counting Kit-8 | YEASEN | Cat No.40203ES76 |
| PrimeSTAR® GXL DNA Polymerase | TaKaRa | Cat No. R050A |
| Fetal bovine serum (FBS) | AMOBIO | Cat No. FBSAB001 |
| Dulbecco’s Modified Eagle Medium | Pricella | Cat No. PM150210 |

| | | |
|---|---|---|
| (RPMI-1640) | Pricella | Cat No. PM150110 |
| **Bacterial Isolates** | | |
| DH5α *E.coli* competent cells | Weidi | Cat. No. DL1004S |
| Rosetta2 (DE3) *E.coli* competent cells | Weidi | Cat. No. EC1014S |
| BL21 (DE3) competent cells | Weidi | Cat. No. EC1002S |
| **Experimental Models: Cell Lines** | | |
| Hela | ATCC | Cat. No. CCL-2 |
| HEK-293T | ATCC | Cat. No. CRL-3216, |
| PANC-1 | Servicebio | Cat. No. STCC11102P |
| MIA PaCa-2 | Servicebio | Cat. No. STCC11101P |
| **Oligonucleotides** | | |
| shPRMT1#1: GTGTTCCAGTATCTCTGATTA | Sangon | N/A |
| shPRMT1#2: CCGGCAGTACAAAGACTACAA | Sangon | N/A |
| sgPRMT1: CGAGGCCGCGAACTGCATCA | Sangon | N/A |

#### RESOURCE AVAILABILITY

##### Lead contact

##### Materials availability

The materials generated in this study are available under a Material Transfer Agreement.

##### Data and code availability

RNA sequencing data have been deposited in the Gene Expression Omnibus (GEO). The accession numbers are provided in the key resources table. The Digital Object Identifier (DOI) is also included in the key resources table. Additionally, all mass spectrometry raw files have been submitted to ProteomeXchange via the PRIDE database. Accessions numbers for these files are also listed in the key resources table.

#### EXPERIMENTAL MODEL AND STUDY PARTICIPANT DETAILS

##### Cell Culture

HEK293T and HeLa cells were obtained from the American Type Culture Collection (ATCC), while PANC-1 and MIA PaCa-2 cells were purchased from Servicebio, China. HEK293T, HeLa, and MiaPaCa-2 cells were grown in Dulbecco's Modified Eagle Medium (DMEM) supplemented with 10% (v/v) fetal bovine serum and 1% penicillin/streptomycin at 37°C in a 5% $CO_2$ atmosphere. In contrast, PANC-1 cells were cultured in RPMI 1640 Medium, also supplemented with 10% (v/v) fetal bovine serum and 1% penicillin/streptomycin, under the same conditions of 37°C and 5% $CO_2$.

##### Animal Studies

Female BALB/c nude mice (5-6 weeks old) were purchased from GemPharmatech, China, and maintained in the specific pathogen-free (SPF) facility of Xiamen University's Animal Laboratory. The mice were housed on a 12-hour light/dark cycle with unrestricted access to standard chow and water. All animal experiments received approval from the Xiamen Animal Care and Use Committee.

#### METHOD DETAILS

##### Protein purification

The cDNA sequences encoding human PRMT1 (residues 32-371) and its mutant proteins were subcloned into a pET-28a vector containing an N-terminal His6-Sumo-StrepI tag. A TEV protease cleavage site was inserted between the Sumo-StrepI tag and PRMT1. Both wild-type (WT) and mutant His6-Sumo-StrepI-TEV-PRMT1 proteins were transformed into *E. coli* BL21 (DE3) cells and subsequently expressed. The cells were cultivated in LB medium at 37°C until an optical density (OD600) of 0.8 was reached, followed by induction with 0.5 mM IPTG at 20°C for 15 hours. For protein purification, the cells were harvested and lysed by sonication in a lysis buffer containing

50 mM Tris-HCl (pH 8.0), 500 mM NaCl, 10% glycerol, and 5 mM 2-mercaptoethanol, supplemented with 1 mM PMSF. After centrifugation, the supernatant was applied to Ni-NTA resin, and the bound protein was extensively washed with Ni-NTA wash buffer (50 mM Tris, pH 8.0; 500 mM NaCl; 10% glycerol; 20 mM imidazole; 5 mM 2-mercaptoethanol) and subsequently eluted using wash buffer supplemented with 250 mM imidazole. Strep-Tactin resin was subsequently employed for further purification after the His6-Sumo tag was cleaved from StrepI-TEV-PRMT1 by SUMO protease. The protein eluted from the Strep-Tactin resin was pooled, concentrated, and loaded onto a Superdex™ 200 Increase 10/300GL size exclusion chromatography column (Cytiva) in a buffer containing 20 mM Tris-HCl (pH 8.0), 150 mM NaCl, and 2 mM DTT. The peak fractions were pooled, concentrated, flash-frozen in liquid nitrogen, and stored at -80°C.

**Negative stain sample preparation**

Wild-type PRMT1, purified via size exclusion chromatography, was prepared at concentrations of 0.1 mg/mL, 0.2 mg/mL, 0.4 mg/mL, and 0.6 mg/mL to investigate the concentration threshold of forming helical polymer. All various concentration samples were prepared following this protocol: First, 5 µL of protein samples were placed on a glow-discharged grid coated with continuous carbon film (Beijing Zhongjingkeyi Technology Co., Ltd.) and incubated for 90 seconds. Second, the excess protein solution was removed using filter paper. Next, the grid was stained with 5 µL of 0.75% (w/v) uranyl acetate for 60 seconds, after which the excess uranyl acetate was removed using filter paper. Finally, the processed samples were examined using a 120 kV electron microscope (FEI Company). All the following mutant PRMT1 samples were prepared at the concentration threshold and observed in negative-stain microscopy with the same protocol.

**Cryo-EM Sample Preparation**

4 µL of 0.6 mg/mL PRMT1 solution was dripped onto the glow-discharged 300 mesh Quantifoil Cu R 1.3/1.2 grid that placed in the Vitrobot (Thermo Fisher Scientific).

The Vitrobot chamber was preset at a temperature of 4 °C and a humidity level of 100%, and the rest parameters were set as following to achieve the desired ice thickness: waiting time of 10 seconds, blotting time of 4 seconds, and blotting force of 1. Then, the grid was promptly plunged into liquid ethane that had been pre-cooled with liquid nitrogen.

**Cryo-EM Data Acquisition**

All the data were acquired using the Titan Krios G3i 300 kV electron microscope in which a spherical aberration corrector, a Gatan continuum (1069) EELS Energy filter and a Post-GIF Gatan K3 Summit direct electron detector are equipped. Program SerialEM[1] is responsible for data collection under super-resolution mode at magnification of 130,000x with pixel size 0.46 Å. The defocus values were set from 1.5 to 2.5 μm. Each movie has 32 dose-fractionated frames with a total electron dose of 50 $e^{-}/Å^{2}$ and an exposure time of 2.09 seconds. 1,748 movies were collected.

**Data Processing**

A total of 1,748 movies were acquired and were firstly processed by MotionCorr2[2] was employed to correct the beam-induced motion of aligning the individual frames, resulting in dose-weighted micrographs with a final pixel size of 0.92 Å. The subsequent data processing was carried out using RELION 3.1.0[3] and lab-made programs. Defocus values for each micrograph were estimated using CTFFIND4[4]. Subsequently, filaments were picked manually from the selected 485 micrographs and high signal-to-noise ratio (SNR) filament segments were kept through 2D (two-dimensional) classification. These high-quality 2D segment classes were employed in the subsequent 3D reconstruction and refinement, with the initial model generated using the lab-made program mrcReference. In addition, we determined the initial helical parameters of helical rise of 19.23 Å and twist angle of 102.86° using our lab-made program called Picker from these data.

Automated filament picking was completed by combining the Topaz program[5], and our lab-made program helixPick. Shortly, all the filament segments were found by the Topaz program and the filament were identified by helixPick from the segment's coordinates. Then, Relion extracted all the filaments following the routine procedure. A total of 2,109,227 segments were extracted and filtered by 2D classification procedure. The selected segments were further used in the subsequent several rounds of Refine3D and three-dimensional (3D) classification and refinement. A final set of 150,914 segments were selected. The map was reconstructed at a resolution of 3.68 Å with the helical parameters, including a twist angle of 104.156 and a helical rise of 24.5843 Å.

**Analytical Gel Filtration**

The size-exclusion chromatography (SEC) analysis of various oligomeric states of human PRMT1, specifically the wild-type PRMT1 oligomer, the Y280A/H296A/T327A mutant PRMT1 dimer, and the W215A/Y220A/F222A mutant PRMT1 monomer, were conducted using a Superdex$^{TM}$ 200 Increase 10/300GL column in SEC buffer (20 mM Tris, pH 8.0; 150 mM NaCl; 2 mM DTT). Thyroglobulin (660 kDa), MLL1 complex (179 kDa), and FTO (55 kDa) served as calibration standards. Protein samples were injected at a flow rate of 0.35 mL/min, and the elution profile was monitored at 280 nm.

**PRMT1 Oligomerization Co-IP Assay**

HEK293T cells (ATCC) were used in PRMT1 oligomerization assay. PRMT1 cDNA (wild-type or mutants) was subcloned into the pCDHEF1 vector with an N-terminal FLAG tag or Myc tag to generate FLAG-PRMT1 or Myc-PRMT1, respectively. Constructs for expressing wild-type or mutant FLAG-PRMT1 and Myc-PRMT1 were co-transfected into HEK293T cells. After 48 hours, the cells were harvested, and the nuclear extracts and cytosolic fractions were prepared following standard protocols. The nuclear extracts and cytosolic fractions were captured using Anti-FLAG resin. Non-specifically bound species were removed by extensive washing with pull-down buffer (50 mM Tris-HCl, pH 8.0; 150 mM NaCl; 10% glycerol; 0.1% NP40; and 2 mM DTT), and bound proteins were eluted using FLAG peptide at 4°C.

Anti-FLAG and anti-Myc signals were examined by Western blotting with the respective anti-epitope antibodies.

**GST/FLAG Pull-down Assay**

Briefly, GST-tagged FUS, hnRNPK, GAR, Fibrillarin (1-83), and Mettl14 (400-456), or FLAG-tagged hnRNPA1 and hnRNPA2, were pre-incubated with GST beads or anti-FLAG resin, respectively, and then mixed with wild-type (WT) or mutant PRMT1 purified from *E. coli* in the binding buffer (50 mM Tris-HCl, pH 8.0; 150 mM NaCl; 2 mM DTT; 10% glycerol; and 0.1% NP40) for 1 hour at 4 °C. After extensive washing with the same binding buffer, the supernatants were eluted using binding buffer containing GSH or FLAG peptide and were subsequently subjected to Coomassie staining.

**CRISPR-Mediated PRMT1 Knockout in Hela Cells**

The PRMT1 knockout (KO) HeLa cell lines were generated using the CRISPR/Cas9 gene-editing system. A single-guide RNA (sgRNA) targeting sequence (5′-CGAGGCCGCGAACTGCATCA-3′) was designed using an online tool (http://crispr.mit.edu) and cloned into the LentiCRISPR v2 vector (Addgene, #52961). HEK293T cells were co-transfected with the LentiCRISPR v2-PRMT1 (gRNA) vector, along with the packaging plasmid psPAX2 (Addgene, #12260) and the envelope plasmid pCMV-VSV-G (Addgene, #8454) using PEI reagent. After 48 hours of transfection, the virus-containing medium was collected and used to infect HeLa cells. The PRMT1-KO HeLa cell line was established through selection with puromycin, and single-cell colonies were isolated. The knockout efficiency was assessed by Western blot analysis and PCR amplification of genomic DNA, followed by sequencing.

**Lentivirus Production and Delivery**

Lentiviruses were packaged for pLKO.1-Hygro-shPRMT1#1, pLKO.1-Hygro-shPRMT1#2, and the control pLKO.1-Hygro-shNC using the packaging plasmid psPAX2 (Addgene, #12260) and the envelope plasmid pCMV-VSV-G (Addgene, #8454). Briefly, 2.25 μg of psPAX2, 0.75 μg of pCMV-VSV-G, and 3 μg of the corresponding constructs were co-transfected into HEK293T cells using PEI (Yeasen Biotechnology, #40815ES03). After 48 hours of transfection, the virus-containing medium was collected and used to infect PANC-1 or MIA PaCa-2 cells at a 1:1 dilution. Forty-eight hours post-infection, 2 mg/mL of Hygromycin B was added to the culture to select for PRMT1 knockdown cell lines. PRMT1 Knockdown efficiency was assessed using Western blot analysis.

**Co-Immunoprecipitation and Arginine Methylation Assay in HEK293T Cells**

Briefly, 3×Flag-tagged full-length hnRNPA1, hnRNPA2, hnRNPK, METTL14, GAR1, FUS, Nucleolin, and Fibrillarin were cloned into the pCDHEF1 vector and co-transfected with various oligomeric states of PRMT1, specifically wild-type Myc-PRMT1$^{oligomers}$, mutant Myc-PRMT1$^{dimers}$, or Myc-PRMT1$^{monomers}$ into HEK293T cells. The nuclear and cytoplasmic extracts were prepared for anti-Flag resin binding for 2 hours. The bound proteins were washed extensively with pull-down buffer (50 mM Tris, pH 8.0, 150 mM NaCl, 10% glycerol, and 0.1% NP40). Subsequently, the proteins were eluted using pull-down buffer containing Flag peptide for Western blot analysis.

**In Vitro Arginine Methylation Assays**

For the PRMT1 enzymatic activity assay, various concentrations of purified wild-type PRMT1 or mutant PRMT1 were incubated with 0.125 μg of histone H4 and 32 μM SAM in 20 μL of methyltransferase assay buffer [50 mM Tris (pH 8.0), 100 mM NaCl, 2.5 mM $MgCl_2$, 1 mM EDTA, and 2.5 mM DTT] at 37 °C for 1 hour. The reaction was terminated by adding 10 μL of 4 × SDS sample loading dye and boiling at 85 °C for 5 minutes. Histone H4 asymmetrically dimethylated at arginine was separated by SDS-PAGE and detected by Western blotting using an anti-H4R3me2a primary antibody.

**Cell proliferation and colony formation assay**

To investigate the role of various oligomeric states of PRMT1 in promoting pancreatic cancer cell growth, shPRMT1-resistant cDNA for PRMT1 was designed and subcloned into the pCDH-EF1-3×FLAG vector. Additionally, site-specific mutations were introduced into this PRMT1 cDNA to produce both dimeric and monomeric forms of PRMT1. Subsequently, the different oligomeric states of pCDH-EF1-3×FLAG-PRMT1 (wild-type or mutant) were transfected into pancreatic cancer cell lines, including PANC-1 and MIA PaCa-2, which are deficient in PRMT1. This ensured that the expression levels of wild-type PRMT1 and its mutants were comparable to those of endogenous PRMT1. The established cell lines expressing distinct oligomeric states of PRMT1, along with untreated pancreatic cancer cells and PRMT1 knockdown pancreatic cancer cells, were utilized for cell proliferation and colony formation assays.

For the cell proliferation assays, the designated cell lines were seeded in 96-well plates at a density of 1,500 cells per well. Proliferation was assessed using the Cell Counting Kit-8. Briefly, the reagent solution was added to each well and incubated at 37°C for one hour at various time points. Subsequently, cell proliferation was quantified by measuring absorbance at 490 nm, using wells containing only medium as blanks.

In the colony formation assay, cells were seeded in 35-mm dishes at a density of 1,000 cells per dish. After approximately 10 days of continuous culture, the cells were washed twice with PBS and subsequently stained with 0.05% crystal violet. The cells were then air-dried and photographed for further analysis.

**Tumor xenograft assay**

The PRMT1-knockdown PANC-1 cell line was generated using the pLKO.1-Hygro-shPRMT1#1 vector. To establish PANC-1 cells that stably express the different oligomeric states of PRMT1, lentiviruses carrying either wild-type or mutant pCDH-EF1-Puro-3×Flag-PRMT1 (oligomer, dimer, and monomer) were packaged and used to infect the PRMT1-knockdown PANC-1 cell line. For the xenograft experiments, female BALB/c nude mice (aged 5-6 weeks) served as recipients. The established cells were harvested, counted, and resuspended at a concentration of 1 million cells per 100 μL in PBS. This cell suspension was mixed in a 1:1 ratio with Matrigel (Corning, 356234), and a total volume of 200 μL was injected subcutaneously into the right flank of the immunocompromised mice. The experimental groups included PANC-1, PANC-1-shPRMT1, and PANC-1-shPRMT1 rescue with PRMT1 oligomer, PRMT1 dimer, or

PRMT1 monomer, all exhibiting expression levels comparable to endogenous PRMT1. Tumor size was measured every 2 days over a total period of 21 days. After the 21-day implantation period, the tumors were harvested, and their weights were recorded. Tumor growth was monitored using calipers, and tumor volume (TV) was calculated using the standard formula: (length × width$^2$)/2.

**Immunoprecipitation LC-MS/MS assay.**

Different oligomeric states of PRMT1 cDNA were subcloned into pCDH-EF1-Puro-3×Flag-PRMT1, along with the pCDH-EF1-Puro-3×Flag-empty vector, for lentiviral packaging to infect PRMT1-depleted PANC-1 cell lines. The resulting cell lines were grown in 15 cm dishes and lysed using a cell lysis buffer containing 10 mM Tris-HCl (pH 7.5), 150 mM NaCl, 1% Triton X-100, 5 mM EDTA, and a complete protease and phosphatase inhibitor cocktail. Lysates were then centrifuged at 4°C for 20 minutes at 13,500 g, and the supernatants were collected. Subsequently, the supernatants were combined with anti-Flag resin for 2 hours of binding. After binding, the resin was collected and washed three times with pull-down buffer (50 mM Tris, pH 8.0, 150 mM NaCl, 10% glycerol, 0.1% NP40), followed by boiling the resin with 1× SDS loading dye for sample preparation. The resulting protein sample were separated using SDS-PAGE, and the gel was cut for subsequent LC-MS/MS analysis.

**RNA sequencing**

Total RNA was extracted from the following cell lines: PANC1, PANC1-shPRMT1, and PANC1-shPRMT1 rescued with PRMT1 oligomer, PRMT1 dimer, and PRMT1 monomer. The expression levels of PRMT1 in these cell lines were comparable to the endogenous PRMT1 levels in untreated PANC1 cells. RNA extraction was performed using Trizol according to the manufacturer's instructions. The construction of the RNA library and sequencing were carried out by Sangon Biotech in Shanghai. Differentially expressed genes (DEGs) were identified using the following cut-off criteria: $|\text{Log2 (fold change)}| \geq 1.5$ and P value $< 0.05$.

**QUANTIFICATION AND STATISTICAL ANALYSIS**

Statistical analysis was conducted using GraphPad Prism 8. For comparisons among more than two groups, one-way ANOVA followed by Tukey's multiple comparison tests was employed. In cases involving two independent variables, two-way ANOVA with Tukey's multiple comparison test was utilized. All statistical analyses were performed using two-tailed tests. Plotted values are expressed as the mean ± SD. P-values below 0.05 were considered statistically significant (*$p < 0.05$, ** $p < 0.01$, ***$p < 0.001$). Pathway enrichment was assessed using Gene Ontology (GO) and Kyoto Encyclopedia of Genes and Genomes (KEGG) tools in Hiplot Pro (https://hiplot.com.cn/) and Reactome gene sets in Metascape (https://metascape.org/).

#### Figure Legends

**Figure S1. Size exclusion chromatography (SEC) elution profile of PRMT1.**

The purified recombinant human PRMT1 protein was analyzed using Superose™ 6 Increase 10/300GL, and the corresponding gel filtration fraction peaks were collected and analyzed using SDS-PAGE.

**Figure S2. Flowchart of wild-type PRMT1 data processing using RELION.**

A total of 1,748 movie stacks were acquired using a cryo-electron microscope at 130,000x magnification. MotionCorr2 (Ref) was employed to align each movie stack, followed by defocus estimation for each micrograph using CTFFIND4. Helix identification and particle picking were performed with the heliPick program based on the particle coordinates provided by Topaz software. Multiple rounds of 2D classification, 3D classification, and Refine3D were conducted to select higher-quality particles and determine more accurate helix parameters. Ultimately, 150,914 particles were selected, resulting in a 3D map with a resolution of 3.68 Å, a final twist angle of 104.156°, and a helical rise of 2.5843 Å.

**Figure S3. The different oligomeric states of PRMT1.**

(A) The size exclusion chromatography (SEC) elution profile of PRMT1 includes wild-type oligomers, Y280A/H296A/T327A mutant dimers, and W215A/Y220A/F222A mutant monomers, which were analyzed using Superdex™ 200 Increase 10/300GL under the same experimental conditions. In this analysis, the corresponding gel filtration fraction peaks were collected and analyzed using SDS-PAGE.

(B) The size exclusion chromatography (SEC) elution profile demonstrates that the Y280A single mutant PRMT1 and the H296A single mutant PRMT1 exhibit a similar elution profile to the Y280A/H296A/T327A mutant PRMT1 dimers. An SDS-PAGE gel image is provided.

(C) Negative staining images and 2D classification of wild-type and mutant PRMT1. W215/Y220/F/222 on the interface between two monomers of dimer were mutated, and PRMT1 cannot form dimer and cannot assemble into helical polymers. 2D classes from these negative stain data explicitly confirm this point. Y280A/H296A/T327A on the interface of two dimers was mutated; PRMT1 can still form dimers but cannot build helical polymers. Their 2D classes further prove this conclusion.

(D) A flowchart illustrating the experimental procedure for the two-tag co-immunoprecipitation assay is provided. Equal amounts of wild-type or mutant FLAG-PRMT1 and MYC-PRMT1 plasmids were co-transfected into HEK293T cells.

(E-F) The nuclear and cytoplasmic extracts from cells in (D) were separately bound to anti-FLAG resin. MYC-PRMT1 binding was assessed using anti-MYC antibodies.

**Figure S4. The helical polymer serves as a preferred structural scaffold for PRMTs binding to RGG motif substrates.**

(A) The domain structure of the hnRNPA1 protein.

(B) In vitro pull-down experiments were conducted. The PRMT1 protein was expressed and purified from *E. coli* to compare its binding affinities with hnRNPA1(1-320)-FLAG, hnRNPA1(1-196)-FLAG, and hnRNPA1(186-320)-FLAG.

(C-D) The size exclusion chromatography (SEC) elution profile of the PRMT1-Fibrillarin complex. The PRMT1-Fibrillarin complex, along with wild-type PRMT1 oligomers and Y280A/H296A/T327A mutant PRMT1 dimers, was analyzed using

Superose™ 6 Increase 10/300GL under consistent experimental conditions. In this analysis, PRMT1 oligomers and PRMT1 dimers served as controls, and the corresponding gel filtration fraction peaks were collected and analyzed using SDS-PAGE.

(E) The cryo-electron microscopy micrograph and the 2D classification results of the PRMT1- Fibrillarin complex are presented.

(F) The 3D reconstruction of the PRMT1- Fibrillarin complex reveals that it is a hexamer.

(G-I) In vitro pull-down experiments were conducted. The PRMT1, PRMT3, PRMT5, PRMT7, and PRMT8 proteins were expressed and purified from *E. coli* or Sf9 insect cells to compare their binding affinities with different RGG motif proteins, including hnRNPA1-FLAG, MBP-Fibrillarin-FLAG, GST-METTL14, and GST-FUS.

**Figure S5. In vitro arginine methylation assays.**

The in vitro methyltransferase activity of wild-type PRMT1 oligomers and mutant PRMT1 dimers against different RGG-rich substrates was evaluated. A series of concentrations (0.2 μM to 12.8 μM) with saturating concentrations of histone H4, hnRNPA1, and fibrillarin as respective substrates were tested to assess the different activities of PRMT1 oligomers and PRMT1 dimers. The protein concentrations of wild-type PRMT1 oligomers and mutant PRMT1 dimers were compared using SDS-PAGE, and the differences in enzyme activity were analyzed through Western blotting.

**Figure S6. PRMT1 oligomerization is required for the growth of pancreatic ductal adenocarcinoma (PDAC) cells.**

(A) HeLa cells were engineered using PRMT1-targeting sgRNAs to generate a PRMT1 knockout (KO) cell line. The PRMT1-KO HeLa cell line was established through selection with puromycin, and single-cell colonies were isolated. Subsequent Western blot analysis was conducted to assess PRMT1 expression levels, with GAPDH serving as the loading control.

(B) The knockout efficiency of the HeLa cell line from (A) was further assessed by

PCR amplification of genomic DNA, followed by sequencing.

(C) HeLa PRMT1-KO cells were transiently transfected with PRMT1 oligomers, dimers, and monomers to achieve PRMT1 expression levels consistent with those of untreated HeLa cells. The results were visualized using Western blot analysis, with GAPDH serving as the loading control.

(D) Cell lysates from (C) underwent gel filtration chromatography to verify the endogenous oligomeric states of PRMT1, with visualization conducted via Western blot.

(E) Western blot analysis detected levels of ADMA, MMA, and H4R3me2a in MIAPACA-2 cells following PRMT1 depletion, with GAPDH included as the loading control.

(F-G) Colony formation and proliferation assays were conducted in PRMT1-deficient cells, followed by analysis using one-way ANOVA.

**Figure S7. Disruption of PRMT1 oligomerization impairs tumor growth.**

(A) Re-expression of oligomeric PRMT1, dimeric PRMT1, or monomeric PRMT1 in MIAPACA-2 cells following PRMT1 depletion was conducted, and Western blot analysis was performed to measure the levels of ADMA, MMA, and H4R3me2a in the cells, using GAPDH as the loading control.

(B-C) The results of colony formation and cell proliferation assays in MIAPACA-2 cells expressing different oligomeric states of PRMT1, along with untreated MIAPACA-2 cells and PRMT1-depleted MIAPACA-2 cells serving as controls, were analyzed using one-way ANOVA.

(D) Immunofluorescence staining of PRMT1 in tumor tissue sections from (main Figure 7J), with a scale bar of 50 μm.

(E) Hematoxylin and eosin (H&E) staining of tumor tissue sections from (main Figure 7J), with a scale bar of 100 μm.

### Supplementary Figure 1. (Ru et al.)

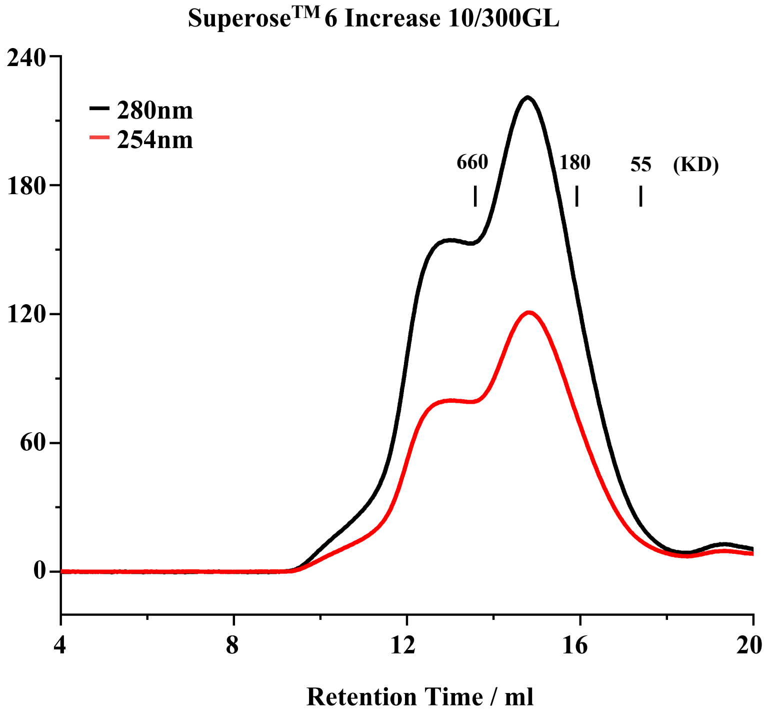

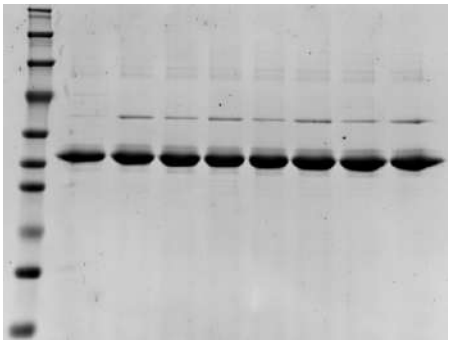

**Supplementary Figure 2. (Ru et al.)**

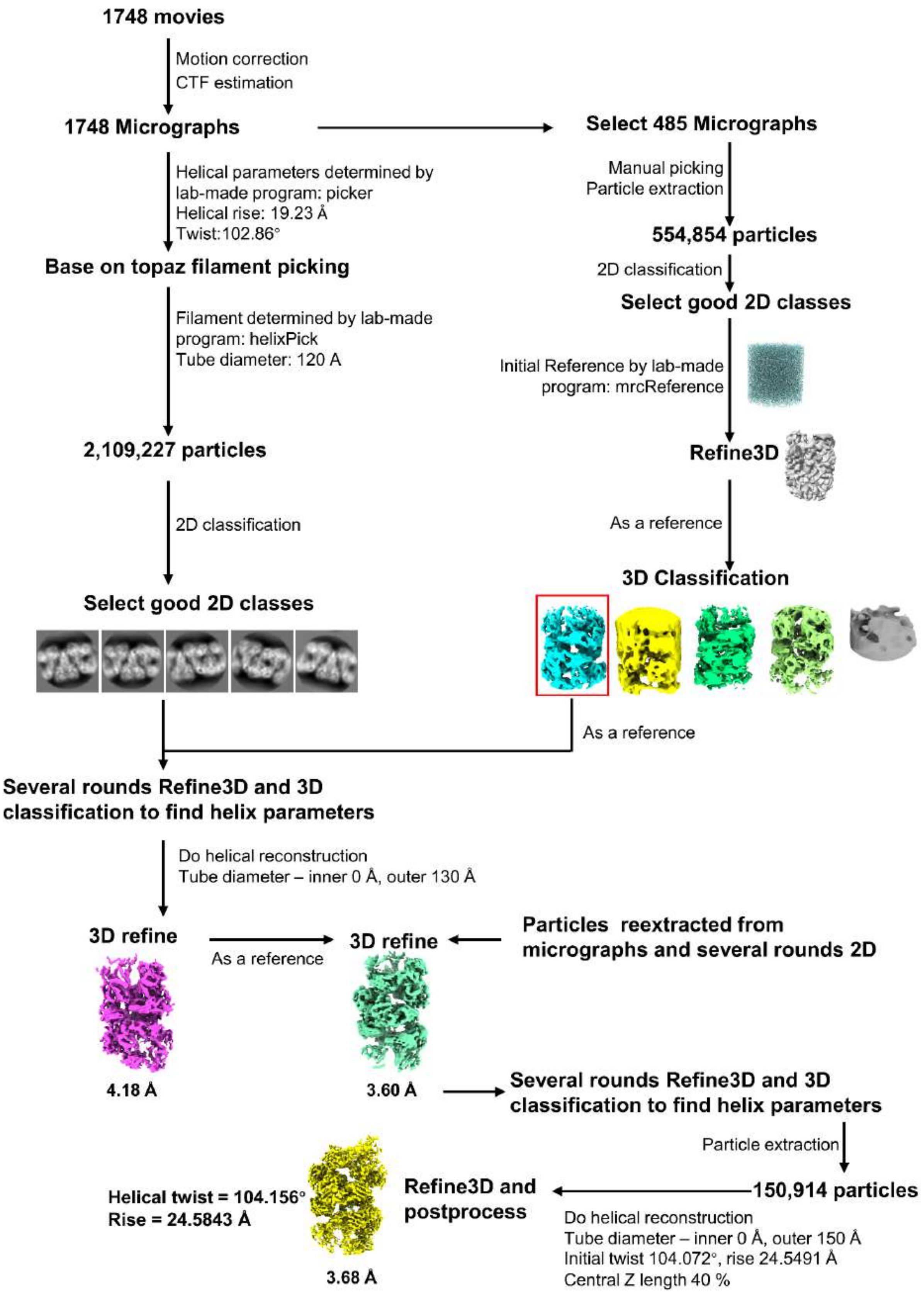

### Supplementary Figure 3. (Ru et al.)

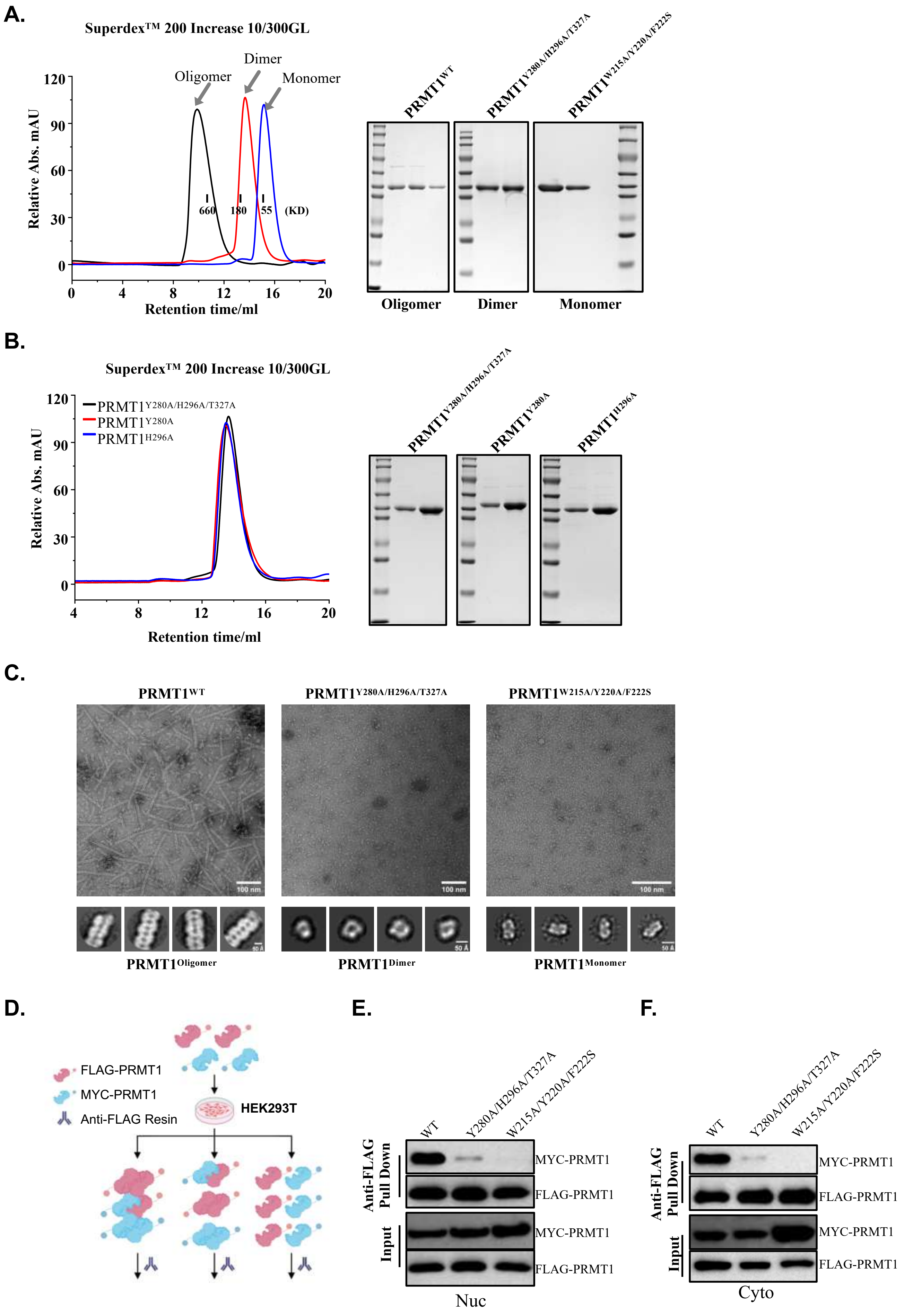

#### Supplementary Figure 4. (Ru et al.)

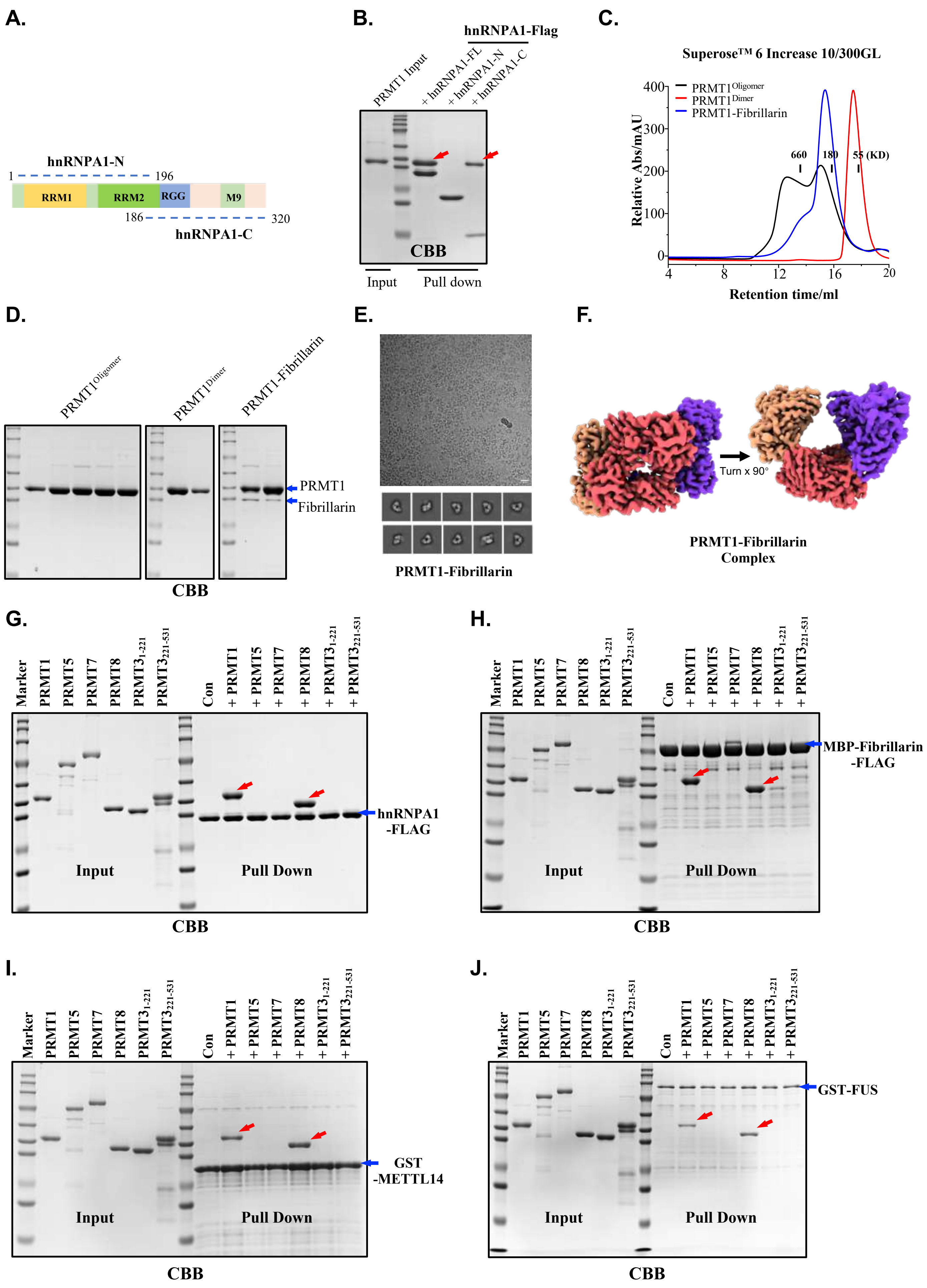

#### Supplementary Figure 5. (Ru et al.)

A.

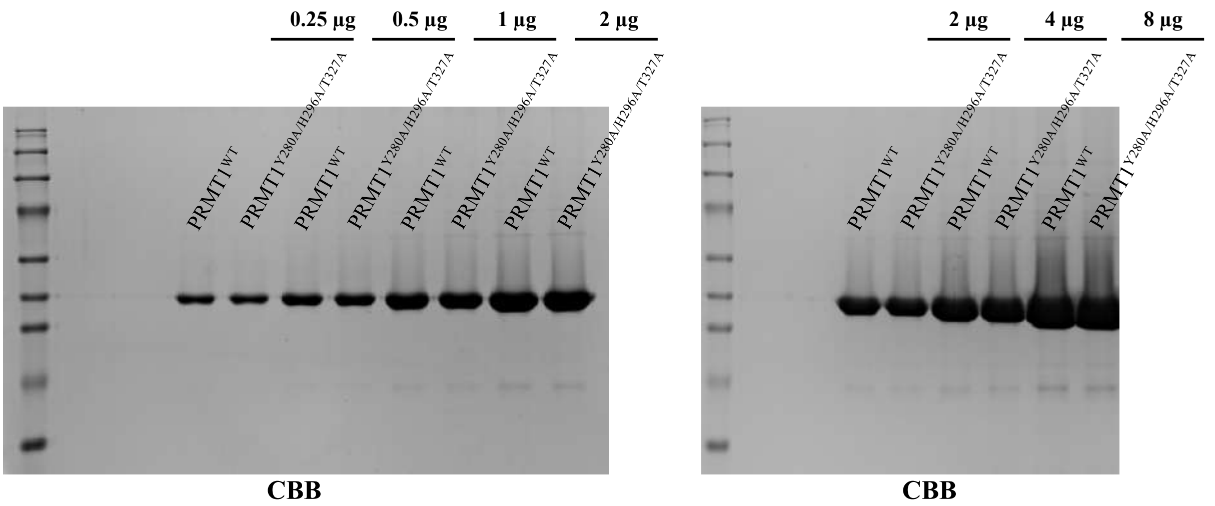

B.

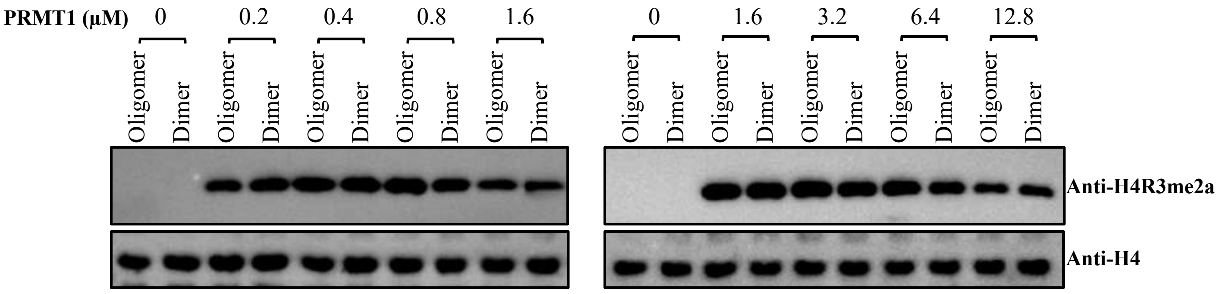

C.

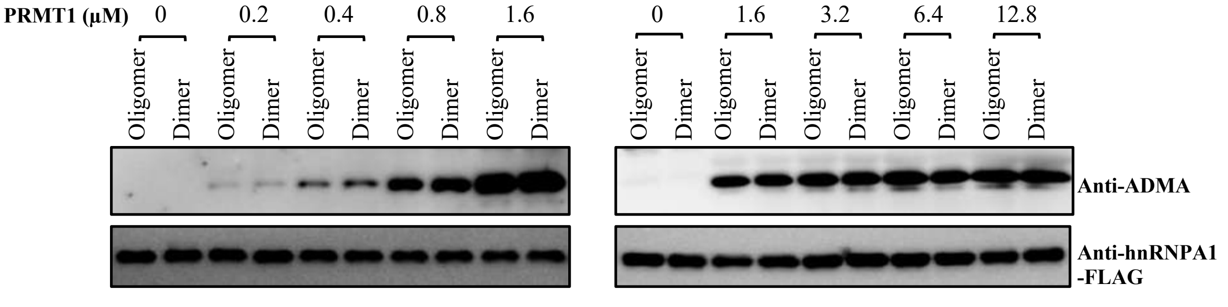

D.

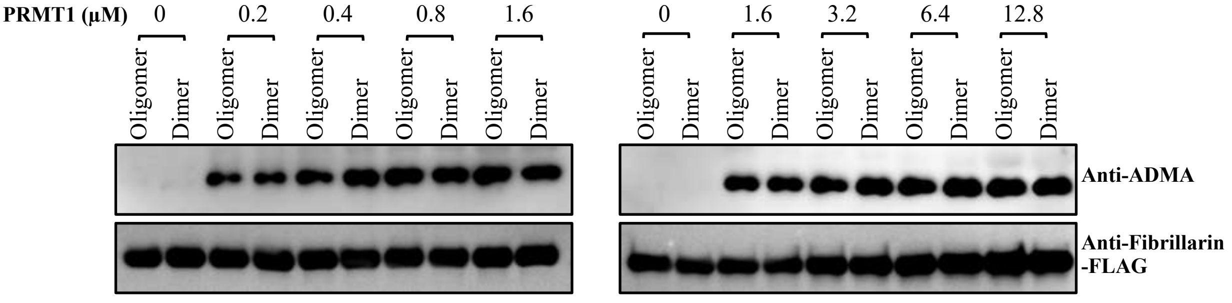

### Supplementary Figure 6. (Ru et al.)

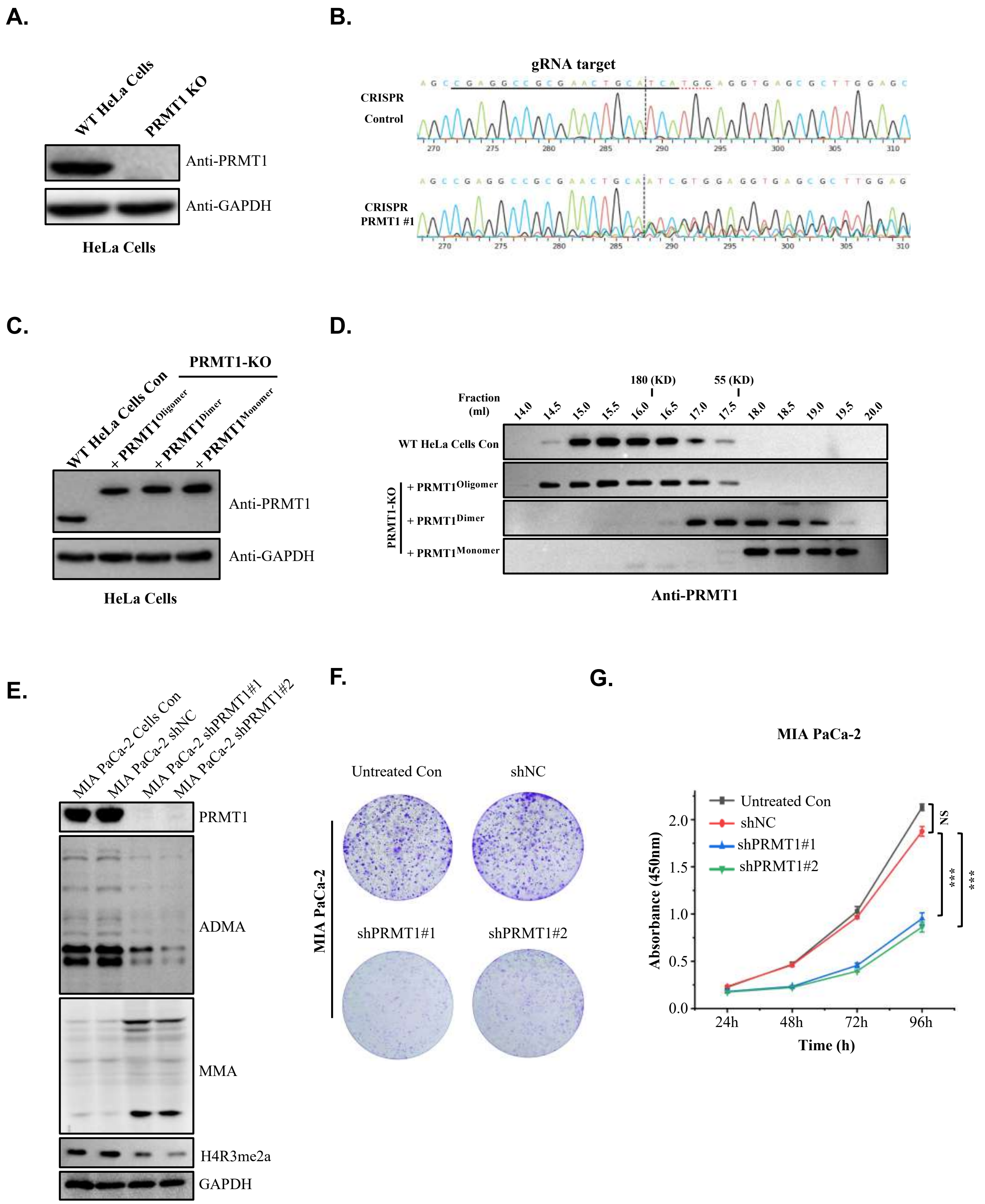

### Supplementary Figure 7. (Ru et al.)

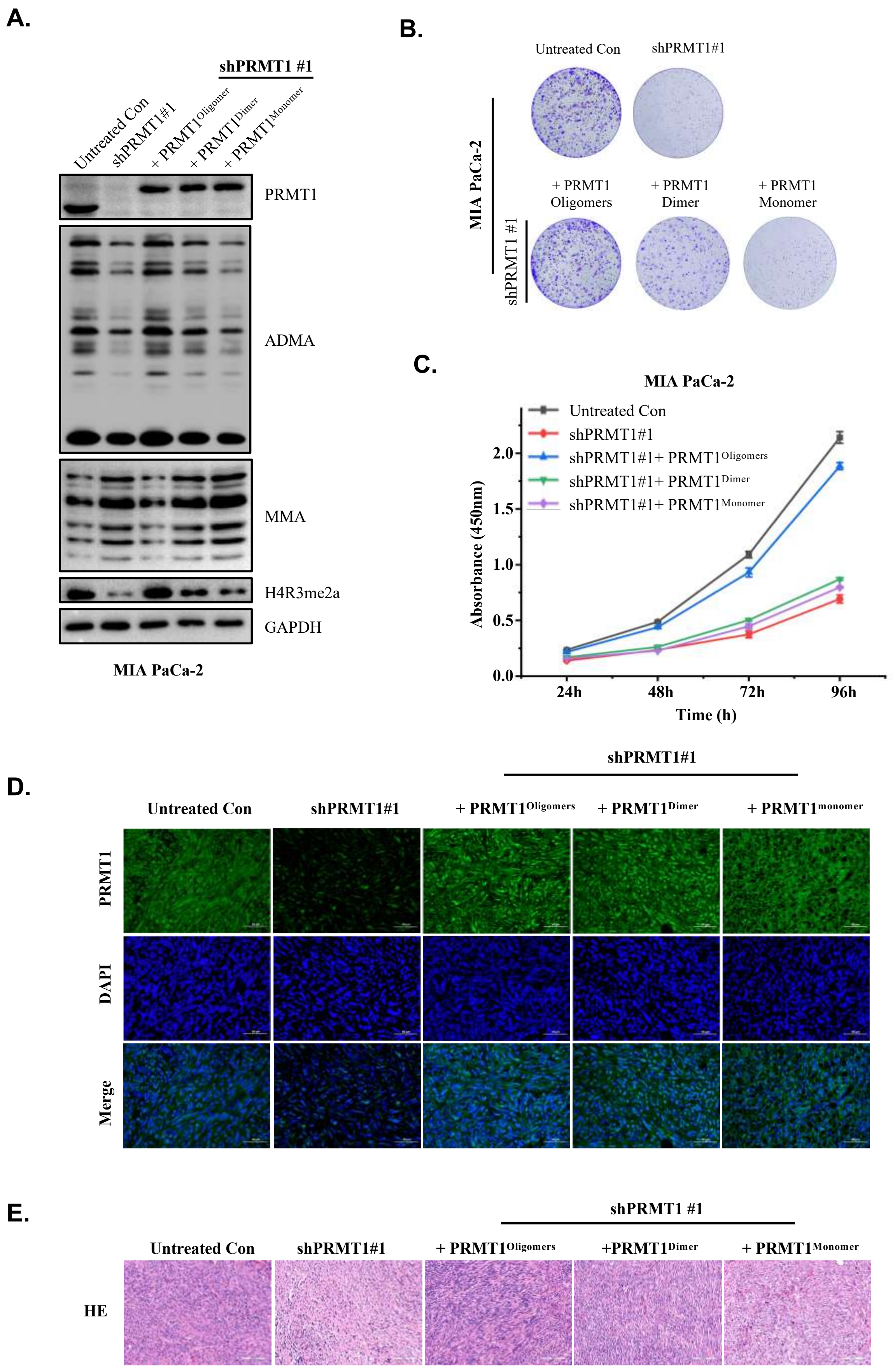

#### Table S1. Cryo-EM data collection, refinement and validation statistics

| | PRMT1<br>(EMDB-61145)<br>(PDB 9J5L) |
|---|---|
| **Data collection and processing** | |
| Magnification | 130K |
| Voltage (kV) | 300 |
| Flux (e-/pix/sce) | 20 |
| Frames per exposure | 32 |
| Electron exposure (e–/Å$^2$) | 50 |
| Pixel size (Å) | 0.92 |
| Micrographs collected | 1,675 |
| Final particle images (no.) | 150,914 |
| Resolution at 0.143 FSC (Å) | 3.68 Å |
| **Refinement** | |
| Initial model used | AlphaFold2 prediction |
| Model resolution (Å) | 3.5 |
| FSC threshold | 0.143 |
| Map sharpening *B* factor (Å$^2$) | -150 |
| Model composition | |
| Non-hydrogen atoms | 25,322 |
| Protein residues | 3,120 |
| Ligands | 0 |
| *B* factors (Å$^2$) | |
| Protein | 154.11 |
| Ligand | 0 |
| R.m.s. deviations | |
| Bond lengths (Å) | 0.004 |
| Bond angles (°) | 0.645 |
| Validation | |
| MolProbity score | 1.90 |
| Clashscore | 9.34 |
| Poor rotamers (%) | 0 |
| Ramachandran plot | |
| Favored (%) | 94.00 |
| Allowed (%) | 6.00 |
| Disallowed (%) | 0 |
